# Sub-diffraction error mapping for localization microscopy images

**DOI:** 10.1101/2021.02.10.427128

**Authors:** Richard J. Marsh, Ishan Costello, Mark-Alexander Gorey, Donghan Ma, Fang Huang, Mathias Gautel, Maddy Parsons, Susan Cox

## Abstract

Assessing the quality of localization microscopy images is highly challenging due to difficulty in reliably detecting errors in experimental data, with artificial sharpening being a particularly common failure mode of the technique. Here we use Haar wavelet kernel analysis (HAWK), a localization microscopy data analysis method which is known to give results without artificial sharpening, to generate a reference image. This enables the mapping and quantification of this common artefact. By suppressing intensity information, we are able to map sharpening errors in a way which is not influenced by nonlinearity in the localisation imaging process. The HAWK Method for the Assessment of Nanoscopy (HAWKMAN) is a general approach which allows the reliability of localization information to be assessed.

---

Image assessment and validation are critical for all microscopy methods, to ensure that the image produced accurately reflects the structure of the sample. When image processing is an integral part of the technique, as in single molecule localization microscopy (SMLM) methods [1, 2], it is important to verify that the image processing is not producing errors or biases. A number of different types of artifact can be introduced by SMLM image processing, including missing structure and artificial sharpening. Artificial sharpening occurs when emitters overlap in the raw data and are incorrectly localized towards their mutual centre, introducing a bias that is often substantial when compared with the estimated localization precision [3, 4, 5, 6, 7].

The importance of assessing the quality of SMLM images is widely recognised but is a challenging problem to solve. In general, images must be assessed without access to the ground truth structure, meaning that any image assessment method must make some type of comparison with alternative data or analysis. For example, Fourier Ring Correlation (FRC) [8] splits the dataset into two images, both of which will be subject to the same algorithmic bias. Therefore, when artificial sharpening is present, FRC will simply report the reduced scatter of localizations typical of biased reconstructions as higher resolution. Most localization algorithms suffer from similar artificial sharpening effects, so comparisons made between them have limited effectiveness [5]. Here, we use the term resolution to indicate the *length scale at which resolvable structures are authentically reproduce in the reconstruction*.

This problem of common biases in each image is averted in the super-resolution quantitative image rating and reporting of error locations method (SQUIRREL) [9]. This method downscales the SMLM image and compares to a linear transformation of the widefield image. However, this has two major disadvantages. Firstly, the downscaling eliminates the fine structure in the image, meaning that only differences on or above the scale of the PSF can be quantified (see Supplementary Note 1 and Supplementary Figs. 1 and 2). Secondly, sharpened images will score more highly than accurate reconstructions if they are more linear in intensity (which is likely for a substantial number of algorithms, see Supplementary Fig. 2).

Two factors contribute substantially (and generally) to nonlinearity in SMLM reconstructions, the background fluorescence and the degree of sampling. The background signal often varies substantially across the field of view and can correlate with the structure, whereas most reconstruction algorithms, as well as SQUIRREL’s linear transform, assume it to be constant. With regard to sampling, the intensity in a widefield (where molecules don’t blink) would normally be an accurate reflection of the labelling density. However, in a typical SMLM experiment, emitters can make multiple appearances (with different intensities) or not at all, meaning that the reconstruction intensity is not reliably related to the number of local fluorophores. This limitation can be circumvented if the sum of the acquired localization microscopy frames is used as the widefield reference. This assures the sampling is the same as in the super-resolution measurement. However, this eliminates the ability to detect missing (non sampled) structure in the test image (although under sampling due to overlapping emitters could still in principle be detected.). To avoid these difficulties, we use a second super-resolution (HAWK pre-processed) image as the reference and both images are binarized to remove the dependence on intensity information (see Supplementary Note 1 for a further discussion of these effects).

There are also approaches to quantify how the algorithm used for SMLM can limit accuracy/resolution and introduce bias [5, 10]. While these can demonstrate bias and artificial sharpening when the ground truth structure (or some defining property such as spatially random structure) is known [5, 10], and can be used to assess the relative performance of algorithms, they cannot assess the quality of reconstructed experimental images. Additionally, the relative performance of different algorithms on simulated test data cannot be guaranteed to reproduce the effects observed on real samples containing varied types of structure.

Here we introduce an alternative approach which allows visual quantification of the accuracy of an SMLM algorithm’s reconstruction. Our method uses HAWK [11], a pre-processing step that for any particular algorithm eliminates the artificial sharpening caused by excessive density in the raw data, at the cost of a small decrease in localization precision. This was demonstrated using ThunderSTORM [12], a standard fitting algorithm that can perform single emitter (SE) and multi-emitter (ME) fitting. HAWK has also been demonstrated to decrease bias, artificial sharpening and nonlinearities in intensity when used with other algorithms.

We exploit the accuracy and reliability of HAWK to identify potential artifacts in a localization microscopy image produced without HAWK, and to indicate where HAWK pre-processing has reduced localization precision sufficiently that underlying fine structure could have been made unresolvable. This is achieved by quantifying structural differences between the original image and the HAWK-processed reconstruction as produced using the same algorithm. This measure can be used to map out areas which have artificial sharpening, and to ascribe a confidence level to local regions of the image at the sub-diffraction level. A measure of the local resolution (in the sense defined above of the length scale at which structure is correctly reproduced), can be ascertained by progressively blurring the input images with a Gaussian kernel for longer length scales and repeating the comparison. The length scale at which reasonable agreement between the HAWK-pre-processed and non-HAWK-processed output images in the local region is achieved, indicates the resolution obtained.

HAWKMAN takes as input data a test (super-resolved reconstructed) image and reference (HAWK-pre-processed) image, and a set of length scales over which the performance of the algorithm will be evaluated (Fig. 1). The input images are intensity-flattened to suppress the influence of isolated outlying high intensity points (due to repeated sampling), which frequently occur in SMLM. The flattened images are then blurred with a Gaussian kernel of width equivalent to the current length scale of interest (ranging from a single pixel up to a user-specified maximum). These blurred images are then binarized according to a length scale-specific adaptive threshold. This produces two images for each input: one (the *sharpening map)* is produced from a threshold of roughly 50% of the local maximum, the other (the *structure map)* is produced from a higher threshold (85%) and subsequently skeletonised. It should be noted that the optimum thresholds will vary slightly depending on the local dimensionality of the sample, but these variations are not critically important. A higher threshold may be required if there is only a small intensity difference between background and structure.

**Figure 1:**
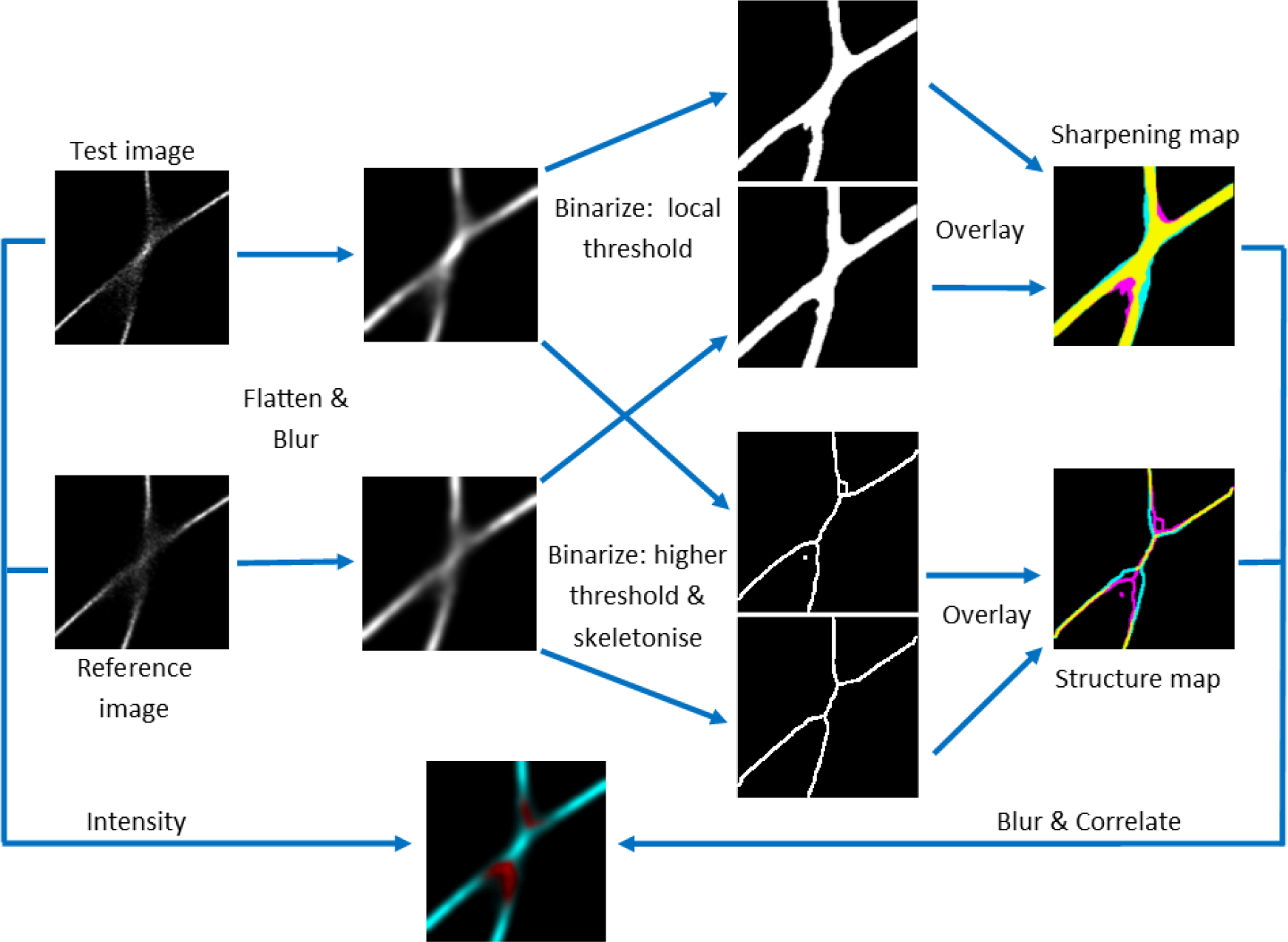
HAWKMAN method and results on experimental data from the Localization Microscopy Challenge. The processing steps of the HAWKMAN algorithm, producing the sharpening map, structure map and confidence map. For the sharpening and structure maps areas of input image agreement are displayed in yellow whereas magenta and cyan indicate structure only present in the test and reference images respectively. Therefore, magenta indicates artificial sharpening and cyan indicates missing structure. The confidence map indicates the calculated local correlation score (see methods for details), ranging from 0 (red) to 1 (cyan), whilst brightness denotes the intensity of the original images.

Differences highlighted in the sharpening and structure maps indicate areas of artificial sharpening, and/or large differences in precision. The local correlation between them is used as a confidence metric for the reliability of that region in the test reconstruction, giving the *confidence map* (Fig. 1). The procedure is repeated for increasing length scales up to the maximum of interest (typically the instrument PSF). This allows the accuracy of the localization algorithm to be assessed across a discrete set of length scales.

In order to test the performance of HAWKMAN we selected simulated datasets from the Localization Microscopy Challenge [13, 14] where both low and high emitter density datasets were available. This allowed us to benchmark the performance of HAWKMAN when using a HAWK reference image, against when using a low density (ground truth) reference image, which ensured that the location and scale of artifacts in the test image was known. Figure 2 shows the results of HAWKMAN analysis on simulations of microtubules for the Localization Microscopy Challenge for both SE and ME Gaussian fitting of the data. For areas of microtubule crossover (Fig.2 a-c) the high density reconstructions (Fig. 2a, b) show substantial sharpening artifacts not present in the reconstruction from low density data (Fig. 2c). As expected, the SE reconstruction (Fig 2a) is more severely affected than the ME (Fig.2b).

**Figure 2:**
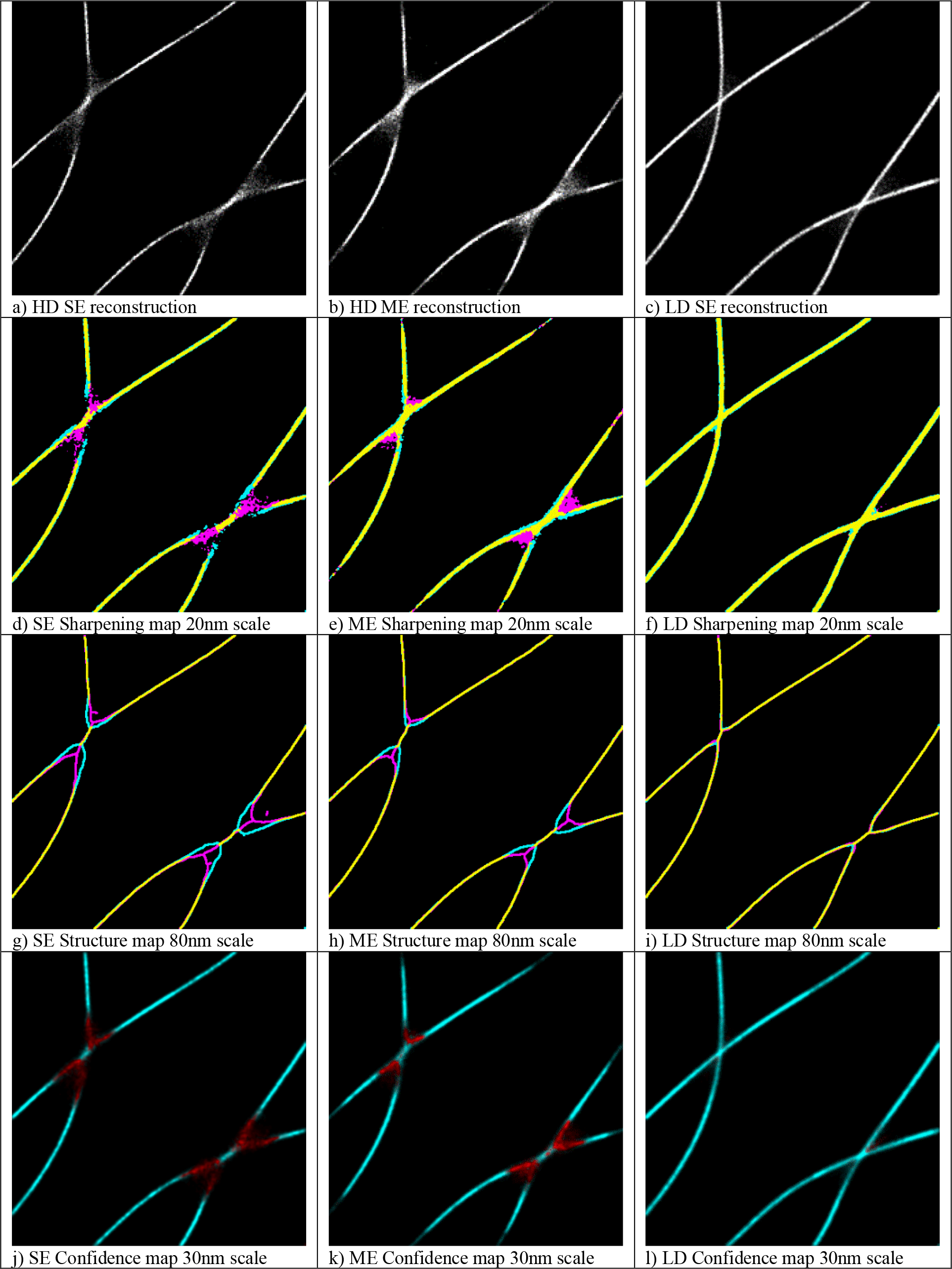
HAWKMAN analysis of simulated microtubule data from the Localization Microscopy Challenge. Comparison of SE and ME fitting of High emitter Density data and equivalent low density data. A section of the reconstructions is shown in (a-c) for HD:SE fitting, HD:ME fitting and LD:SE fitting respectively, which contain artificial sharpening for both high density methods. The results of HAWKMAN analysis are shown in (d-l) for each of the three test reconstructions with HAWK used as the reference. The sharpening maps (d-f) and structure maps (g-i) are shown at a length scale of 20nm and 80nm respectively. In both the SE and ME cases the magenta areas (test image only) in the sharpening maps (d,e) and the structure maps (g,h) indicate substantial sharpening along with missing structure (cyan), which is more severe (as expected) in the SE case. Both these effects are absent for the low density reconstruction (f,i). These results are reflected in the confidence maps (j-l, 30nm scale). These show substantially reduced confidence (strong artifacts, highlighted in red) for both high density methods (j,k), whereas for the low density data the confidence (l) is high everywhere.

HAWKMAN analysis was performed on the challenge data sets, with the reference image being produced by the same algorithm on the same data but with prior HAWK pre-processing. Sharpening maps (Fig. 2d-f), structure maps (Fig.2 g-i) and confidence maps (Fig. 2j-l) are shown for the same length scales for each test image. The length scales in this figure (and all subsequent ones) were selected to give a clear and concise representation of the full HAWKMAN output (see Supplementary Videos 1-5 for the full sets of HAWKMAN output for each result). For clarity of display the structure map has been dilated [15].

For the high density reconstructions, both the sharpening and structure maps show substantial biases in the microtubule positions, indicated by the magenta (test image only) where the microtubules cross. Cyan (reference image only) areas indicate some missing structure for the high density images not present in the low density reconstruction. The structure maps indicate these persist at even quite large (80nm) length scales. This ‘pinching in’ of crossing microtubules is a commonly observed artifact in high density data [4, 7, 9, 11]. The estimated degree of error is quantified in the confidence maps (low confidence marked in red, high confidence in cyan) for both SE (Fig. 2j) and ME (Fig. 2k) methods. These indicate ME fitting is more accurate than the SE fitting, but still produces substantial errors. The corresponding confidence map for the same structure simulated at low emitter density (Fig. 2l) correctly indicates no sharpening artifacts are present at this length scale (30nm)

The Localization Microscopy Challenge datasets also contain experimental data, and we tested the performance on high density experimental microtubule datasets [13, 14]. These were analysed using both SE & ME Gaussian fitting [12], giving results with clear artificial sharpening (Fig. 3 a,b) in the regions of clustering and microtubule crossover. The corresponding reconstruction when HAWK is used (Fig. 3 c,d) show much more clearly resolved structure in these regions, but some precision is lost. The output of HAWKMAN analysis indicates that these are highly sharpened reconstructions. The sharpening maps (Fig. 3 e,f), structure maps (Fig. 3 g,h) and confidence maps (Fig. 3 i,j) indicate the presence of severe sharpening artifacts even at quite large length scales (60nm, 160nm & 100nm respectively).

**Figure 3:**
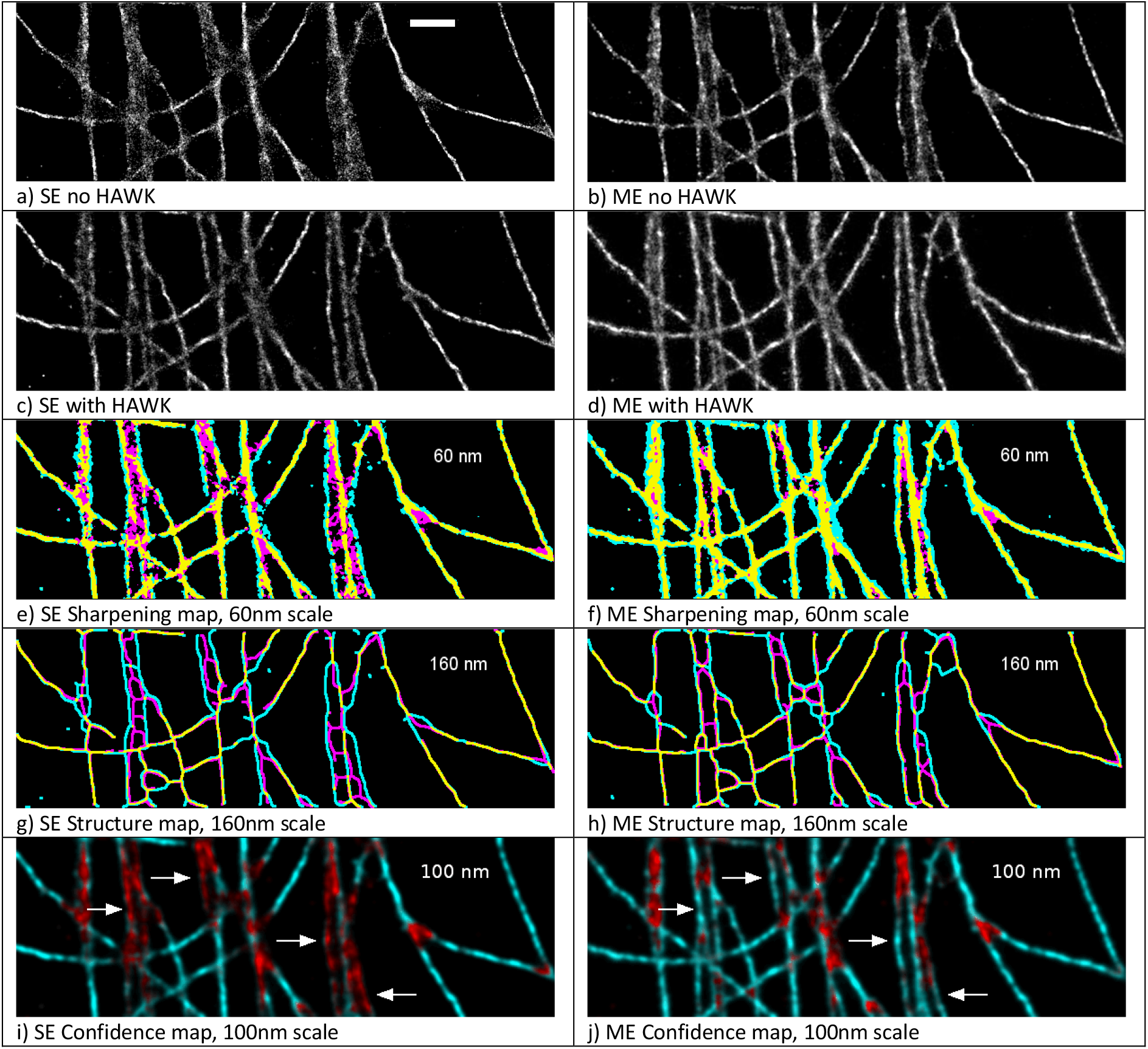
Results of HAWKMAN analysis on experimental microtubule data from the Localization Microscopy Challenge comparing SE and ME fitting. The reconstructions for SE and ME fitting without HAWK are shown in a) and b) respectively along with their equivalents c) and d) when HAWK is used. The results of HAWKMAN analysis are shown in the sharpening maps (e,f), the structure maps (g,h) and the confidence maps (i,j) for selected length scales. The SE reconstruction (a) shows substantial density-induced sharpening artifacts. The differences in structure between this reconstruction and the results of applying HAWK with SE fitting are highlighted by the sharpening map (e) at a scale of 3 pixels (60nm). Substantial bias in the SE reconstruction where microtubules are in close proximity/crossing is highlighted in the structure map (g) even at a scale of 160nm. Reliable regions or the reconstruction where both methods agree are highlighted in the confidence map (i) at a scale 100nm. ME fitting (b) gives a more accurate reconstruction, however, as can be seen in (f,h,j) it still contains numerous artifacts. The confidence map (100nm scale) (j) for this reconstruction shows a reduction in the severity of artifacts but some remain. The white arrows highlight some of the structure accurately reported (at this length scale) in the ME result (j) that were inaccurate in the SE case (i). Scale bar 1 μm.

Examining the reconstruction produced by ThunderSTORM ME fitting [12] (Fig. 3b), it can easily be seen that this is a superior reconstruction to SE fitting (Fig. 3a), but some structures are still not clearly resolved. The confidence map produced by HAWKMAN analysis for this reconstruction is shown in Fig. 3i (100nm scale). The map shows a much higher confidence in the overall accuracy of this reconstruction, yet still highlights some areas where sharpening is present. At other scales, (Supplementary Figs. 3, 4; Supplementary Movies 1, 2) and with other high density analysis algorithms, SRRF & SOFI [16, 17], (Supplementary Fig. 5; Supplementary Movies 3, 4, 5), HAWKMAN clearly demonstrated the presence and scale of artificial sharpening. It should be noted that for some localization algorithms, the output reconstruction can vary significantly with the choice of parameter values used for the analysis [11, 16]. Supplementary Fig. 5 demonstrates, with the example of SRRF, how HAWKMAN may aid in the optimisation of input parameters. The relative fidelity of reconstructions using different parameter sets can be assessed by using HAWKMAN on each of them.

Tests on other simulated data, also from the Localization Microscopy Challenge, demonstrate that HAWKMAN is clearly able to detect and even quantify sharpening and artifacts. This is the case even at length scales substantially below the diffraction limit (see Supplementary Fig. 6), right down to the scale of the localization precision. The advantage of using a HAWK image as the reference is that it does not contain the same bias common to many other algorithms. Using a HAWK-processed high density dataset as a reference image provides comparable results to using a low density (i.e. unsharpened) dataset, in contrast to when other algorithms are used for the reference image (see Supplementary Fig. 7). This confirms the suitability of HAWK processing for a qualitative and quantitative assessment.

The importance of removing intensity information is demonstrated by comparing HAWKMAN with SQUIRREL. Even at the PSF scale HAWKMAN is able to correctly identify the relative fidelity of differently sharpened reconstructions in situations where SQUIRREL cannot (Fig. 4 and Table 1). Here assessment of experimental microtubule data from the Localization Microscopy Challenge is performed using both HAWKMAN and SQUIRREL for comparison. A widefield image produced by summing image frames (Fig. 4a) shows areas where microtubules are poorly resolved (even compared to the widefield - blue arrows) and a high intensity area (yellow arrow). The reconstruction produced by SE and ME fitting are shown in Fig. 4b, where a Gaussian blur of scale equivalent to the PSF has been applied (the scale at which SQUIRREL makes its comparison). Comparing these with the widefield image shows significant differences, particularly in the intensity. The error maps produced by SQUIRREL are displayed below these (Fig. 4c). They show the largest errors are reported where there are large intensity differences between the reconstructions and the widefield reference. For easier comparison with the HAWKMAN results the relative error in the SQUIRREL maps was converted into a confidence score (see Methods) and used to colourise the blurred reconstructions (Fig. 4d) in the same manner as for HAWKMAN. Comparison with the HAWKMAN results at the PSF length scale (Fig. 4e) highlights how SQUIRREL has detected the intensity differences (yellow arrow), but largely ignored the differences in structure (blue arrows). The converse is true for HAWKMAN, which indicates low confidence in areas of structural dissimilarity, and high confidence in the areas that differ only in intensity. HAWKMAN also reports the ME result as a much more authentic reconstruction than that from SE fitting, something that can be confirmed by inspection of the areas marked by blue arrows. Conversely, SQUIRREL rates these similarly, as any errors arising from structural differences are swamped by the much larger intensity error common to both reconstructions.

**Figure 4:**
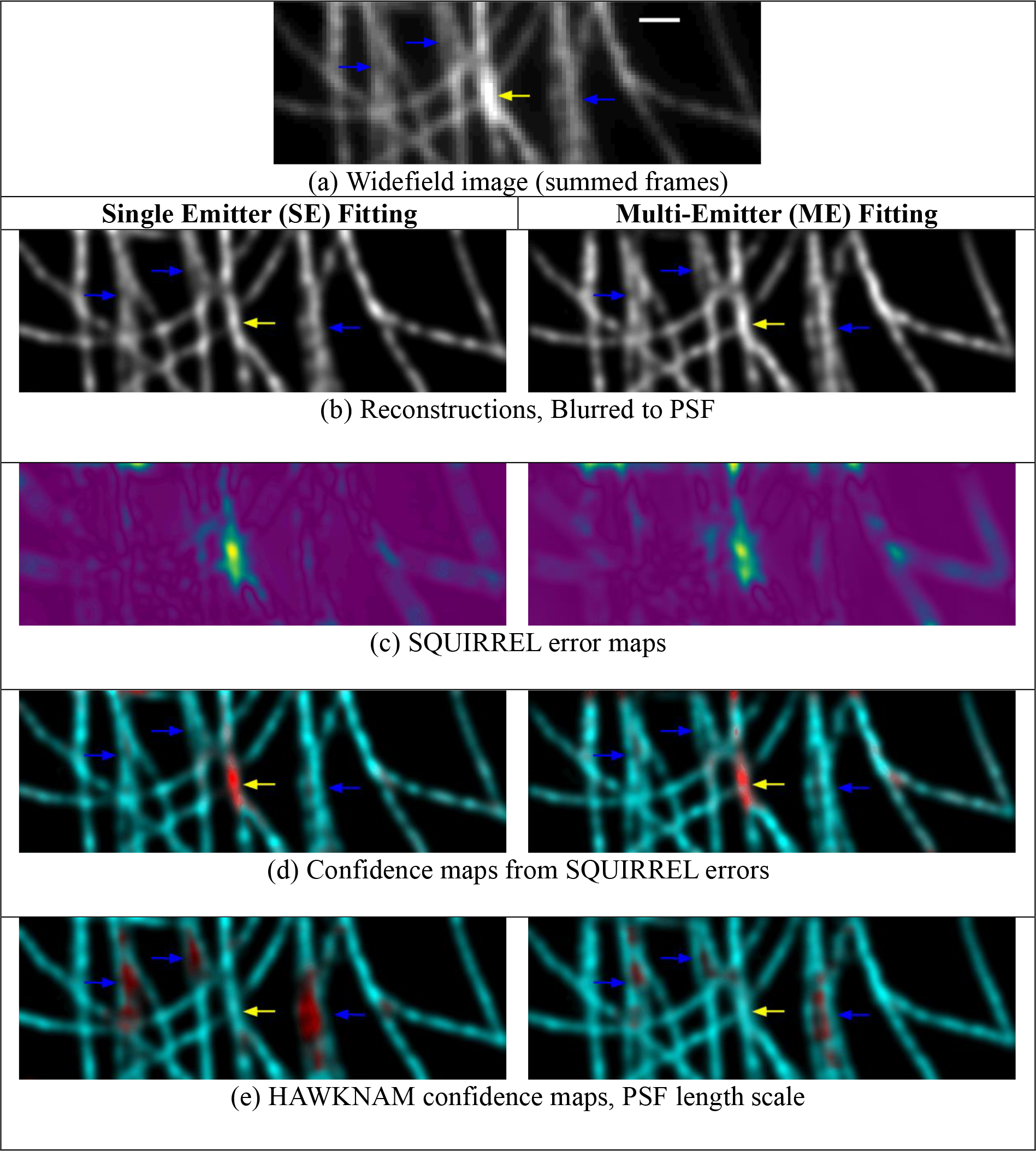
Comparison of the ability of SQUIRREL and HAWKMAN to detect structural artifacts when faced with intensity differences in the input and reference images, on experimental high density data from the Localization Microscopy Challenge. (a) A pseudo widefield image produced by summing image frames. (b) SE and ME reconstructions. Blue arrows indicate areas of obvious sharpening compared to the widefield, yellow arrow denotes an area of high intensity. (c) Error maps produce by SQUIRREL analysis show the largest errors (yellow) are dominated by the high intensity region in (a). The relative error is used instead of the confidence score to produce a HAWKMAN style confidence map (d) from the SQUIRREL data. When compared to the actual HAWKMAN result (e), it shows SQIRREL principally detects the intensity differences and does not recognise the superior structural details (blue arrows) of the ME reconstruction.

Table 1 shows the quality metrics produces by SQUIRREL for the above results. These measure the correlation (Resolution Scaled Pearson Correlation Coefficient) and error (Resolution Scaled Error) between the downscaled super-resolution image and the widefield. These are compared with similar quantifications for HAWKMAN, the correlation coefficients between the binarised test and reference images in the sharpening and structure maps (PCC Sharpening and PCC Structure respectively). These show that SQUIRREL is unable to detect that the ME reconstruction is superior to the SE case, as its errors are dominated by the same intensity difference. The HAWKMAN correlation scores, along with the confidence maps above clearly indicate a much more accurate reconstruction with ME fitting as expected.

**Table 1:**
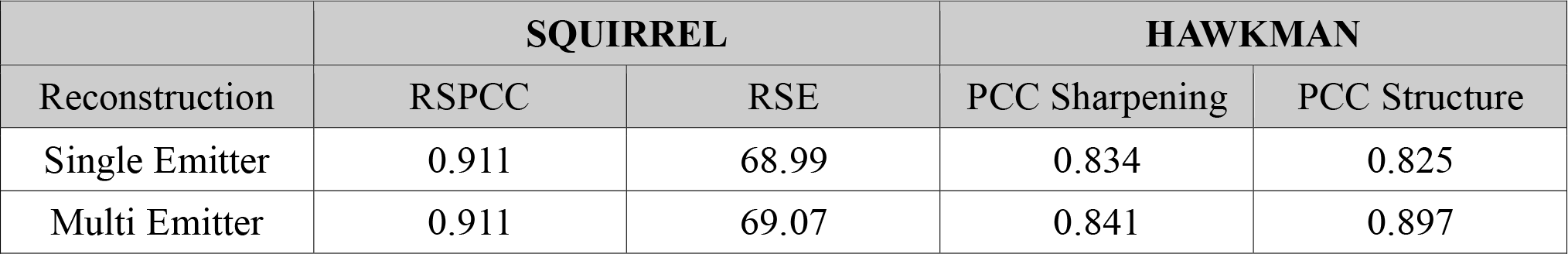
Comparison of reconstruction quality metrics for SQUIRREL and HAWKMAN on Challenge Microtubule data. The table displays the quality metrics output by the two methods on the high density data analysed in Figure 4. For SQUIRREL both the Resolution Scaled Pearson Correlation Coefficient and the Resolution Scaled Error are unable to determine which is the better reconstruction (ME actually has marginally higher error). However, for HAWKMAN both the correlation between sharpening maps and structure maps is significantly higher for the Multi-Emitter result, even at the PSF scale, indicating this is the higher fidelity reconstruction.

| Reconstruction | SQUIRREL |  | HAWKMAN |  |
| --- | --- | --- | --- | --- |
|  | RSPCC | RSE | PCC Sharpening | PCC Structure |
| Single Emitter | 0.911 | 68.99 | 0.834 | 0.825 |
| Multi Emitter | 0.911 | 69.07 | 0.841 | 0.897 |

Using HAWK with many algorithms does introduce a small decrease in localization precision compared to when that algorithm is used alone [11]. The magnitude of which typically increases with the severity of bias present in the reconstruction without HAWK. For any given localization algorithm, as the emitter density increases, the precision of the HAWK reconstruction will be reduced but will remain unsharpened. Although this leads to reduction in the resolution attained in the HAWK image, the improvement in fidelity over the reconstruction without HAWK will increase. This means that when compared with an image of the ground truth (for instance, a low density reconstruction), using the HAWK-reconstruction as the reference image slightly exaggerates differences with the input image. This may lead to a slight over-estimation of errors in some instances.

Additionally, large differences in precision between the HAWK and non-HAWK images may indicate the emitter density has exceeded the capabilities of the algorithm, even when used in combination with HAWK, resulting in reduced resolution. HAWKMAN can indicate where in a reconstruction and at what length scale the precision of the HAWK reference is sufficiently reduced that it may conceal unresolved finer structure. The experimenter is thus warned of this situation and may decide a lower emitter density is required. Alternatively, a higher performance more computationally intensive localization algorithm may be required, such as switching from SE to ME fitting.

A demonstration of how this effect can enable HAWKMAN to detect that there may be missing fine structure, even in the HAWK reference image, is demonstrated using simulation (Supplementary Figs. 8 and 9). For a limited range of emitter densities, structure can be unresolvable but comparable in scale to the resolution in the HAWK image. Thus, even though structure is not visible in the HAWKed or non-HAWKed images, HAWKMAN may be able to give a first order approximation to the underlying structure. This effect is demonstrated on the Z1Z2 domain in sarcomeres imaged at high density (see Supplementary Fig. 10 for simulated data and Supplementary Fig. 11 for experimental data). Here structure is not visible in either input image but is present in the HAWKMAN confidence map at a 50-70nm length scale, which is in line with the expected observation for this structure.

To further asses the performance of HAWKMAN on experimental data we selected three well-known biological structures of differing geometry and scale to demonstrate the performance of HAWKMAN on experimental data: clathrin-coated pits, microtubules and mitochondria. This allowed us to test the performance of the system on small structures a few hundred nanometres in size (Fig. 5 a-d), extended linear structures (Fig. 5 e-h), and structures with an extended fluorophore distribution (Fig. 5 i-l). Differences between SE-fitted data processed without (Fig.5 a, e, i) and with HAWK (Fig. 5 b, f, j) are visually clear and are reflected by the HAWKMAN sharpening and confidence maps. These are displayed at an appropriate length scale for each dataset Fig. 5). As mentioned previously, a modified value of the threshold coefficient may be required for optimum performance when analysing low-contrast structures; this was the case for the clathrin data (see Methods and Supplementary Fig. 12 for details).

**Figure 5:**
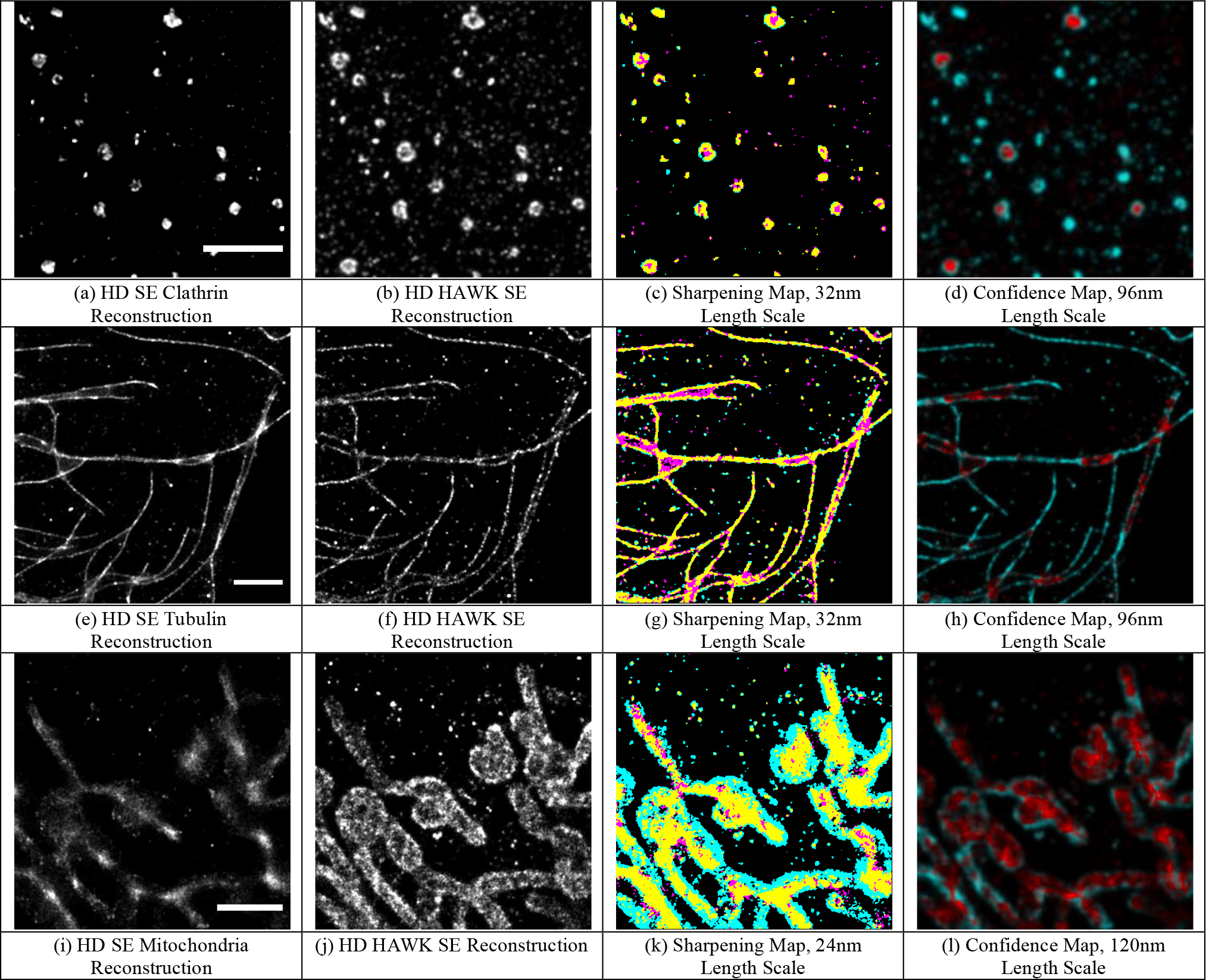
HAWKMAN can assess reconstructions from a variety of different structures. Here, clathrin (a-d), tubulin (e-h), and the mitochondria epitope TOMM20 (i-l) have been imaged at medium-to-high density and evaluated with HAWKMAN. The test images were reconstructed using SE ThunderSTORM (a, e, i), and analysed with a reference image similarly reconstructed from the HAWK pre-processed dataset (b, f, j), and analysed using the sharpening map (c, g, k) and the confidence map (d, h, l). Data is displayed at a length scale of 32nm (c, g) and 24nm (k) for the sharpening maps, and at 96nm (g, h) and 120nm (l) for the confidence maps. The sharpening map threshold coefficients were 0.9 (c) and 0.7 (g, k). The confidence maps (d, h, l) highlight the substantial degrees of sharpening, particularly the TOMM20 reconstructions (i-l). Colouring as in Fig. 1, scale bars 2μm.

Experimentally, artificial sharpening is particularly likely to be experienced in multi-colour dSTORM measurements due to the variable performance of dyes. While Alexa Fluor^®^ 647, with its high-intensity, low density emissions is exceptionally well-suited to dSTORM acquisitions, most other dyes are far less so. Researchers looking to perform multi-colour dSTORM analysis must therefore contend with the fact that dyes whose emission spectra peak in other laser channels blink with much lower brightness and at much higher density, making them highly prone to the generation of sharpened data (a situation also likely to be true when using imaging living cells). HAWKMAN can identify the regions in which artificial sharpening is occurring in typical two-colour dSTORM data (see Supplementary Fig. 13).

The basis of HAWKMAN is that it exploits the fact that algorithms fail in different ways when used with and without HAWK. At extreme activation densities most algorithms become significantly biased, whereas with HAWK there is no bias but a decrease in localization precision. Therefore, when both images agree, one can be confident that the activation density is appropriate for that algorithm and the image contains the most accurate and precise localizations it can produce (i.e. comparable to a low density acquisition).

Image artifacts are a critical issue in localization microscopy. They can be very difficult to detect and quantify, even for experienced users, as they can involve subtle structure changes. In addition, they may only be present in certain regions of the image, with many parts of the image appearing of good quality. Therefore, providing tools which can detect artifacts below the resolution limit is critical if SMLM is to provide reliable and trustworthy results. HAWKMAN is a straightforward, fast (less than 1 minute on a desktop PC for the challenge data presented in Fig. 3) and reliable test, available as an ImageJ [15] plugin, which users can apply to verify the quality of their data.

## Supporting information

Plugin

## Acknowledgements

We thank D. Matthews for his assistance with the N-STORM system in the Nikon Imaging Centre at King’s College London. This work was supported by the BBSRC (BB/R021767/1) and the MRC (MR/R008264/1). SC acknowledges support from a Royal Society University Research Fellowship and a Royal Society Enhancement award and IC acknowledges support from an EPSRC studentship.

## Author Contributions

R.J.M and S.C. conceived and designed the analysis and algorithm, with assistance from M.P., R.J.M. wrote the analysis software and produced simulations. All analysis of Localization Microscopy Challenge data was performed by R.J.M. Preparation of samples and experimental measurements of microtubule and clathrin-coated pit data were performed by A-M.G and I.C. Mitochondria samples and data where prepared and collected by D.M and F.H. Analysis of experimental data was performed by I.C aided by R.J.M and S.C. Myofibril samples were prepared by M.G. and R.J.M. performed the experiments. The manuscript was produced by R.J.M., I.C. And S.C. All authors reviewed the manuscript.

## Data availability statement

The raw image sequences and reconstructions of original data supporting this research will be downloadable from the Kings College London Server on publication. Raw data from the Localization Microscopy Challenge is publicly available from http://bigwww.epfl.ch/smlm/

## Code availability statement

The HAWKMAN analysis software in the form of an ImageJ plugin is provided as Supplementary Software.

## Methods

### HAWKMAN sharpening detection

HAWKMAN analysis enables the similarity of two SMLM images to be assessed in a way which does not rely on the intensity values in the images being linearly related, enabling us to reliably detect bias in the positions of structures. It should be noted that while this method will detect artifacts caused by errors in the SMLM data analysis, but not artifacts which cause fluorophores to be imaged in a different location to the protein of interest (e.g. labelling artifacts). Therefore, to fully assess the authenticity of a super-resolution image, multiple methods of assessment may be necessary.

Two images are compared: the test image (which is undergoing quality assessment) and the reference image (which is produced using HAWK pre-processing and a fitting algorithm such as ThunderSTORM). For validating HAWKMAN, we have also sometimes used fits of low density data for the reference image, as a proxy for the ground truth. To avoid very bright points (from repeated localizations) leading to variable thresholding performance, we cap the maximum pixel intensity to the 98^th^ percentile of the intensity histogram (excluding zeros).

A Gaussian blur is then applied to the image. This blur determines the length scale at which the quality of the data is being assessed. By using multiple sizes of blur, the local resolution (i.e. the length scale at which information starts to become locally unreliable) can be determined (see Supplementary Fig. 6). The algorithm will assess successive blur levels up to a user-specified maximum, with an increment of the pixel size of the super-resolution image. The images are then normalised to a maximum intensity of one. From this starting point, three different mappings of errors are produced, each likely to highlight errors under different conditions.

The first assessment which we calculate is the sharpening map. An adaptive threshold for each image is calculated using Wellner’s method [18]. Here we use a neighbourhood size of the largest odd number of pixels not greater than the current length scale. This produces a map of the mean intensity around each pixel. The images are then binarised based on whether the pixel intensity is above a set proportion *(C_b_*) of the local threshold 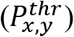.

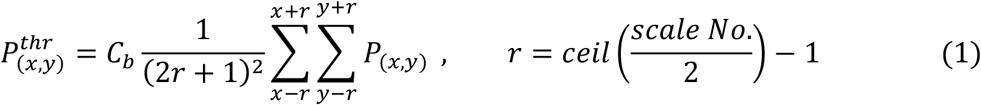

Where *ceil(x)* is the function that raises *x* to the first integer greater than *x*. The value of *C_b_* determines what pixels in the image are classed as structure and which are classed as background. A value of *C_b_* = 0.7 leads to a thresholded image where the width of linear structures is roughly equal to the FWHM of the intensity profile of a line profile through the structure in the original image. This value is designed to pick up features which have a peak-to-trough intensity ratio of about two. Where the image values between parts of a structure does not fall to this level, a higher threshold may be needed. This can be seen in the clathrin-coated pits imaged in Fig. 5 of the main text: here, the depth of the central hole is sometimes shallower than a factor of two, so a higher coefficient of 0.9 was used to also detect sharpening in these shallower pits. For all simulations and experimental data from the Localization Microscopy Challenge [14] and muscle sarcomeres, the standard coefficient of 0.7 was used.

At small blur scales, local variations in labelling density and multiple reappearances of individual fluorophores can lead to falsely identifying these as very fine structure (microtubules are a common example of this). To counter this, we determined experimentally that an additional threshold of 0.1 times the calculated threshold at a neighbourhood size of half the PSF helped smooth out this false fine structure. Additionally, a small baseline contribution to the threshold *(C_a_* = 0.04) eliminated many background localizations and reduced the fixed-pattern noise of some algorithms.

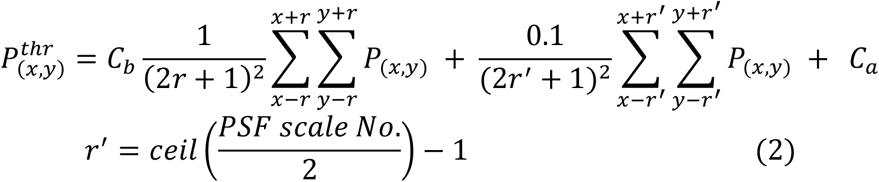

A colour overlay of the binarized images is then produced to reveal sharpening artifacts. Areas where there are substantial differences between the images are indicative of sharpening in the test image or loss of precision in the reference. Areas where the two images agree can be considered reliable.

This ‘sharpening map’ is appropriate for detecting structure thinning, but less suited to visualising the collapse of adjacent structures to one. The binarisation of the blurred and flattened images is repeated but with a higher threshold coefficient *C_b_* = 0.85 (but never lower than the sharpening threshold if this is increased) and a baseline threshold of *C_a_* = 0.02 to detect local maxima. These binarised images are then skeletonised using standard methods [15] giving a skeletonised interpretation of the structure. Again, a composite image of where the two structures agree and disagree reveals differences in structure collapse.

Similarity between the test and reference output images is measured using the two-dimensional cross correlation for both the sharpening and structure images (where the structure images are first Gaussian-blurred to the current length scale so as to become progressively tolerant of differences below this scale). A ‘confidence map’ is produced by summing the normalised, blurred test and reference images. This is then colourised according to the calculated local correlation, which indicates how likely the reconstruction is to be unbiased. Here, any local correlation above 0.85 in both the sharpening and skeletonised images is deemed as indicating valid structure. Areas below this level are colourised according to the level of agreement as measured by this correlation. The confidence level *S_conf_* is given by:

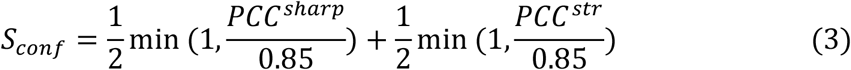

where PCC^sharp^ and PCC^str^ are the Pearson correlation coefficients for the sharpening and structure comparisons respectively, calculated over a range equal to the current length scale, and min(x,y) indicates the smaller of x or y.

The above process is repeated for each length scale, producing an output image sequence wherein the degree of local sharpening is mapped as a function of scale. The actual reliable resolution obtained locally in the reconstruction can be assessed by observing at which scale disagreement between the test and the reference starts to occur.

### Experimental Data

#### Tubulin & Clathrin

##### Fixation & Labelling

Human cervical cancer cells (HeLa) were thawed and cultured in Dulbecco’s Modified Eagle Media (DMEM) 1% Penicillin and Streptomycin (PS, Sigma-P0781), 10% Foetal Bovine Serum (FBS, HyClone-SV30160.03) and 1% L-Glutamine (L-Glu, Sigma-59202C). Cells were transferred to a T25 flask (Cellstar-690175) and incubated at 37°C and 5% Carbon dioxide. Passaging cells were trypsinized with 10% trypsin (Sigma-T4174) diluted in Phosphate-Buffered Saline (PBS, Sigma-D8537), for 5 min once cells reached 80% confluence. Complete DMEM was added to neutralize trypsin and centrifuged at 1200 rpm for 3 min. The supernatant was aspirated, and cells resuspended in 5ml of complete DMEM. Cells were plated on Fibronectin (Sigma-FC010) coated 35mm high glass bottom dishes (ibidi-81158) for imaging Cells were fixed for 15 min in 3.6% Formaldehyde (PeqLab-30201) at room temperature and washed 3 times with PBS.

Permeabilisation and blocking were undertaken by incubating cells in ‘blocking buffer’ consisting in 3% BSA (Sigma-10735108001) & 0.5% Triton X-100 (Sigma-X100) in 1×PBS for 10 minutes. Cells were then incubated with primary antibodies (tubulin from Sigma-T8328 at 1:200, clathrin from Abcam-ab21679 at 1:500) diluted in blocking buffer for 30 minutes while gently rocked at room temperature (RT). After three 5 minute washes in ‘washing buffer’ consisting in 0.2% BSA & 0.1% Triton X-100 in 1×PBS, cells were incubated with the Alexa Fluor^®^ 647 secondary antibodies (Invitrogen-A21235 for tubulin, Invitrogen-A21244 for clathrin, both at 1:500 dilution) in blocking buffer for 30 minutes, gently rocked at RT. Cells underwent three five minute washes in PBS×1 before being stored at 4°C in 1×PBS for up to two days before imaging.

##### Imaging & Hardware

Cells were imaged in imaging buffer, where 1220μl buffer was made using 800μl distilled water, 200μl ‘MEA’, 20μl ‘GLOX’, and 200μl ‘dilution buffer’. Here, ‘MEA’ is 1M cysteamine (Sigma-30070), and 0.25M HCl (Sigma-H9892); ‘GLOX’ is 35mM glucose oxidase (Sigma-G6125), 13.6μM Catalase (Sigma-C40), 8mM Tris (Amresco-E199), and 40mM NaCl; lastly, ‘dilution buffer’ is 50mM Tris, 10mM NaCl (Alfa Aesar-A12313), and 10% w/v glucose (Thermo Fisher-G/0450). Dilution buffer was added just before imaging to minimise cell damage due to pH change.

Objective-based total internal reflection fluorescence (TIRF) was employed to minimise background. Fluorophore bleaching was undertaken with widefield illumination prior to image acquisition to minimise background signal from above the structures of interest. Approximately 5,000 of a total 20,000 10ms frames were acquired with supplementary 405nm activation to obtain high density data.

Experimental data of clathrin and tubulin was gathered from fixed HeLa cells on a Nikon motorized inverted microscope ECLIPSE Ti2-E with Perfect Focus System in the King’s College Nikon Imaging Centre. It is equipped with a laser bank with 405nm, 488nm, 561nm, and 640nm lasers (LU-NV series), a 160nm-pixel ORCA-Flash 4.0 sCMOS (scientific Complementary metal-oxide-semiconductor, Hamamatsu Photonics K.K.), and a CFI SR HP Apochromat TIRF 100XAC oil objective (NA 1.49) with an automatic correction collar.

#### Muscle Sarcomeres

##### Sample preparation

Mouse cardiac myofibrils were prepared from freshly excised mouse cardiac muscle fibre bundles that had been tied to plastic supports to maintain an average sarcomere length of about 2.4 μm, as verified by laser diffraction. The fibre bundles were stored over night at 0°C in rigor buffer (140 mM KCl, 2 mM MgCl2, 1 mM EGTA, 2 mM DTT, 20 mM HEPES, pH 6.8, containing protease-inhibitors (Roche)). The next morning, the central sections of the fibres were dissociated by mechanical dissociation with a homogenizer following the protocol by (Knight, P.J. Meth. Enzymol., 1982. PMID: 7121291), washed in rigor buffer 5 times and stored in rigor buffer and at 0°C until use. Suspensions of myofibrils in rigor-buffer were adhered to poly-lysine coated glass-bottomed dishes and fixed with 4% PFA in rigor buffer, washed in phosphate-buffered saline (PBS) and incubated in PBS/10% normal goat serum, before incubating with the rabbit polyclonal antibody Z1Z2 (binding to an epitope at the N-terminus of titin; Young, P EMBO J., 1998. PMID: 9501083), which labels the myofibrils at the Z-disc. Goat anti-rabbit secondary antibodies conjugated to Atto647N (50185, Sigma) or Alexa647 (A-21244, Life Technologies) were applied after washing the samples in PBS, and visualisation performed after washing away unbound secondary antibody with PBS.

##### Imaging and Hardware

The apparatus used has been described in more detail in [11]. Briefly, this consisted of a customised STORM microscope, built around a DMi8 Microscope body and ‘SuMo’ passively stabilized stage (Leica-microsystems GMBH). In this system the 1.43 160X objective (Leica-microsystems GMBH) is mounted to the underside of the stage via a piezo drive (PI). Focus was maintained using a custom active control system. Excitation was from a 638nm diode laser (Vortran). Emitter density was controlled with a 405m Diode laser (Vortran). Fluorescence was collected in the 660-700nm spectral range using dichroic filters (Croma). The imaging of sarcomere samples was performed in a standard reducing buffer (Glox-glucose,200mM MEA).

#### Mitochondria

##### Fixation and labelling

COS-7 cells (CRL-1651, ATCC) cultured on 25-mm-diameter coverslips (CSHP-No1.5-25, Bioscience Tools) were first fixed with 37 °C pre-warmed 3% PFA (15710, Electron Microscopy Sciences) and 0.5% GA (16019, Electron Microscopy Sciences) in 1×PBS (10010023, Gibco) at room temperature (RT) for 15 min. Subsequently, cells were washed twice with 1×PBS and then quenched with freshly prepared 0.1% NaBH4 (452882, Sigma-Aldrich) in 1×PBS for 7 min. After washing three times with 1×PBS, cells were treated with 3% BSA (001-000-162, Jackson ImmunoResearch) and 0.2% Triton X-100 (X100, Sigma-Aldrich) in 1×PBS for 1 h, gently rocked at RT. Then cells were incubated with primary antibodies (sc-11415, Santa Cruz Biotechnology) at 4 °C overnight. After washing three times for 5 min each time with wash buffer (0.05% Triton X-100 in 1×PBS), cells were incubated with secondary antibodies (A21245, Molecular Probes) at RT for 5 h. Both primary and secondary antibodies were diluted to 1:500 in 1% BSA and 0.2% Triton X-100 in 1×PBS. After washing three times (5 min each time with wash buffer), cells were post-fixed with 4% PFA in 1×PBS for 10 min. Then cells were washed three times with 1×PBS and stored in 1×PBS at 4 °C until imaging.

##### Imaging buffers and sample mounting

Immediately before imaging, the coverslip with cells on top of it was placed on a custom-made holder. 40 μL of imaging buffer (10% (w/v) glucose in 50 mM Tris (JT4109, Avantor), 50 mM NaCl (S271-500, Fisher Chemical), 10 mM MEA (M6500, Sigma-Aldrich), 50 mM BME (M3148, Sigma-Aldrich), 2 mM COT (138924, Sigma-Aldrich), 2.5 mM PCA (37580, Sigma-Aldrich), and 50 nM PCD (P8279, Sigma-Aldrich), pH 8.0) was added on top of the coverslip. Then another coverslip was placed on top of the imaging buffer. This coverslip sandwich was then sealed with two-component silicon dental glue (picodent twinsil speed 22, Dental-Produktions und Vertriebs GmbH).

##### SMLM Setup

SMLM imaging was performed on a custom-built setup on an Olympus IX-73 microscope stand (IX-73, Olympus America) equipped with a 100x/1.35-NA silicone-oil-immersion objective lens (FV-U2B714, Olympus America) and a PIFOC objective positioner (ND72Z2LAQ, Physik Instrumente). Samples were excited by a 642 nm laser (2RU-VFL-P-2000-642-B1R, MPB Communications), which passed through an acousto-optic tuneable filter (AOTFnC-400.650-TN, AA Opto-electronic) for power modulation. The excitation light was focused to the pupil plane of the objective lens after passing through a filter cube holding a quadband dichroic mirror (Di03-R405/488/561/635-t1, Semrock). The fluorescent signal was magnified by relay lenses arranged in a 4*f* alignment to a final magnification of ~54, and then split with a 50/50 non-polarizing beam splitter (BS016, Thorlabs). The split fluorescent signals were delivered by two mirrors onto a 90° specialty mirror (47-005, Edmund Optics), axially separated by 580 nm in the sample plane, and then projected on an sCMOS camera (Orca-Flash4.0v3, Hamamatsu) with an effective pixel size of 120 nm. A bandpass filter (FF01-731/137-25, Semrock) was placed just before the camera.

#### Simulations

Simulated data for parallel lines were taken from data used in the original evaluation of HAWK (see [11] for detailed methods). Briefly, MATLAB was used to generate simulated microscopy image sequences using a Gaussian PSF of 270nm and a camera pixel size of 100nm. The labelling density was 100/μm along the lines. The active emitter density was controlled by varying the ‘off’ time. The number of frames was varied, so that in each case emitters make an average of five appearances regardless of the emitter density. Realistic photon and camera noise were included.

For the simulation of Z1Z2 antibody labelling of sarcomeres these simulations were reperformed with the following modifications. Instead of each line consisting of a single column of fluorophores, three adjacent columns on the 10nm spaced grid contained fluorophores, simulated a line of width 20nm. The spacing between line centres was 60nm resulting in an observable gap of 40nm. The emitter density was controlled by varying the emitter off time until the output closely resembled the experimental result. This corresponded to an activation density of 33 emitters/μm^2^ of structure. Note this results in a high degree of emitter overlap in each frame than the same activation rate on the previous simulations due to the higher labelling density. The degree of sampling of each emitter was reduced to an average of 0.75 appearances per emitter, again this was adjusted for the best subjective match to experiment.

#### Data Analysis

All acquisitions underwent ThunderSTORM analysis as performed using the single and ME methods. Settings were as follows: Filter - Difference of Gaussians (min 1.0, max 1.6 pixels), Detector method - local maximum (connectivity 8), PSF model - Integrated Gaussian, Estimator - Maximum likelihood, fitting radius = 3. For ME fitting additional/alternate parameters were: fitting radius = 5, max molecules = 5, p-threshold = 0.05. SOFI analysis was used using the publish MATLAB script [https://www.epfl.ch/labs/lben/lob/page-155720-en-html/page-155721-en-html/] which performs the calculation to 4th order. In all cases the balanced output was used. SRRF analysis was performed using the ImageJ plugin [https://doi.org/10.1038/ncomms12471] using the default parameters, except for where HAWKMAN was used to optimise some values, as described in Supplementary Fig 5. Compressed sensing was performed using the published MATLAB script [https://doi.org/10.1038/nmeth.1978]. The model PSF was set to match that used in the simulations. DeconSTORM was also performed using MATLAB [http://zhuang.harvard.edu/decon_storm.html] with input parameters for fluorophore blinking were set to match the simulations. Due to computational limitations the convolution method was used over the matrix method. Where HAWK pre-processing was used, identical parameters for the subsequent localization step were used as in the case where HAWK was not used. The only additional processing was the filtering of false positive localizations by fitted width, following the procedure described in the original publication [11].

##### Simulated line pairs

Image sequences where analysed using ThunderSTORM [12] (single and ME fitting), Balanced SOFI [17], SRRF [16], compressed sensing (CSSTORM) [19] and DeconSTORM [20]. For ThunderSTORM, SOFI and SRRF the full-size image of 64×64 pixels (100nm size, 270nm PSF) was analysed. Due to the large computational requirements of CSSTORM and DeconSTORM, the sequence was first cropped to 17×17 pixels for these methods. The reconstructions from the other algorithms where subsequently cropped to the equivalent area. Any offset between the different methods was adjusted at this stage, the required offset having been established by simulating a single point emitter. For all methods except SOFI, a magnification factor of 10 was used, giving a reconstruction pixel size of 10nm. For SOFI, the maximum magnification factor was 4, so these images where converted to 10nm pixels size using bilinear interpolation (ImageJ).

##### Localization Microscopy Challenge: simulated data

For all analysis algorithms the same methods and settings where used as above, making allowances for the differences in camera pixel size and gain. DeconSTORM was not used for this dataset, as the blinking properties of the simulated emitters were not known. Only ThunderSTORM was used on the low density simulation, as no significant emitter overlap should be present in this data.

##### Localization Microscopy Challenge: experimental microtubule data

For both SE and ME fitting with ThunderSTORM, all parameters were as above, except a magnification factor of 5 was used to render the images (corresponding to a 20nm reconstruction pixel size). For SRRF with ‘default parameters’: ring radius = 0.3, Axes in ring = 6, Temporal analysis = TRPPM. For SRRF using ‘optimum parameters’: Ring radius = 0.3, Axis in ring = 8, Temporal analysis = TRAC order 2. In both cases the magnification factor was 5.

##### Sarcomere data

This data was analysed using ThunderSTORM SE fitting using the same parameters as above. The magnification was x10 (10nm pixel size).

##### Clathrin-coated pit, mitochondria, and microtubule data

These data sets were all analysed using ThunderSTORM SE fitting, using the same parameters as the Localization Microscopy Challenge data, with the exception of the differing camera pixel sizes these data sets where acquired on (160nm for the microtubules and clathrin, 120nm for TOM20). As the mitochondrial data was acquired on a bi-focal plane microscope, only a single plane was analysed.

##### HAWKMAN analysis

For all the analysis presented here, default threshold parameters were used, except for the clathrin-coated pit data, for which the thresholds were raised to 0.9 as described above and in the main text. The reconstructions using the default SRRF parameters on the Localization Microscopy Challenge data included a low intensity fixed-pattern noise background of ~5% of the maximum intensity. To prevent analysing this as structure, this background was subtracted from the images before processing. This stage was not necessary for all other data.

## References

[1] E. Betzig, G. H. Patterson, R. Sougrat, O. W. Lindwasser, S. Olenych, J. S. Bonifacino, M. W. Davidson, J. Lippincott-Schwartz and H. F. Hess, “Imaging intracellular fluorescent proteins at nanometer resolution,” Science, pp. 1462–5, 2006.

[2] M. J. Rust, M. Bates and X. Zhuang, “Stochastic optical reconstruction microscopy (STORM) provides sub-diffraction-limit image resolution,” Nature Methods, vol. 3, no. 10, pp. 793–5, 2006.

[3] S. Wolter, U. Endesfelder, S. v. d. Linde, M. Heilemann and M. Sauer, “Measuring localization performance of super-resolution algorithms on very active samples,” Optics Express, vol. 19, no. 8, pp. 7020–33, 2011.

[4] P Fox-Roberts, R. Marsh, K. Pfisterer, A. Jayo, M. Parsons and S. Cox, “Local dimensionality determines imaging speed in localization microscopy,” Nature Communications, 2017.

[5] E. A. K. Cohen, A. V Abraham, S. Ramakrishnan and R. J. Ober, “Resolution limit of image analysis algorithms,” Nature Communications, 2019.

[6] S. v. d. Linde, S. Wolter, M. Heilemann and M. Sauer, “The effect of photoswitching kinetics and labeling densities on super-resolution fluorescence imaging,” Journal of Biotechnology, vol. 149, no. 4, pp. 260–6, 2010.

[7] A. Burgert, S. Letschert, S. Doose and M. Sauer, “Artifacts in single-molecule localization microscopy,” Histochemistry and Cell Biology, vol. 144, no. 2, pp. 123–31, 2015.

[8] R. P. J. Nieuwenhuizen, K. A. Lidke, M. Bates, D. L. Puig, D. Grünwald, S. Stallinga and B. Rieger, “Measuring image resolution in optical nanoscopy,” Nature Methods, vol. 10, pp. 557–62, 2013.

[9] S. Culley, D. Albrecht, C. Jacobs, P. M. Pereira, C. Leterrier, J. Mercer and R. Henriques, “NanoJ-SQUIRREL: quantitative mapping and minimisation of super-resolution optical imaging artefacts,” Nature Methods, p. 263–6, 2018.

[10] S. Mailfert, J. Touvier, L. Benyoussef, R. Fabre, A. Rabaoui, M.-C. Blache, Y Hamon, S. Brustlein, S. Monneret, D. Marguet and N. Bertaux, “A Theoretical High-Density Nanoscopy Study Leads to the Design of UNLOC, a Parameter-free Algorithm,” Biophysical Journal, vol. 115, no. 3, pp. 565–76, 2018.

[11] R. J. Marsh, K. Pfisterer, P Bennett, L. M. Hirvonen, M. Gautel, G. E. Jones and S. Cox, “Artifact-free high-density localization microscopy analysis,” Nature Methods, p. 689–692, 2018.

[12] M. Ovesný, P Křížek, J. Borkovec, Z. Švindrych and G. M. Hagen, “ThunderSTORM: a comprehensive ImageJ plug-in for PALM and STORM data analysis and super-resolution imaging,” Bioinformatics, p. 2389–90, 2014.

[13] D. Sage, H. Babcock, T.-A. Pham, T. Lukes, T. Pengo, J. Chao, R. Velmurugan, A. Herbert, A. Agrawal, S. Colabrese, A. Wheeler, A. Archetti, B. Rieger, R. Ober, G. M. Hagen, J.-B. Sibarita, J. Ries, R. Henriques, M. Unser and S. Holden, “Super-resolution fight club: assessment of 2D and 3D single-molecule localization microscopy software,” Nature Methods, vol. 16, no. 5, pp. 387–95, 2019.

[14] D. Sage, H. Kirshner, T. Pengo, N. Stuurman, J. Min, S. Manley and M. Unser, “Quantitative evaluation of software packages for single-molecule localization microscopy,” Nature Methods, vol. 12, no. 8, pp. 717–29, 2015.

[15] C. A. Schneider, W. S. Rasband and K. W. Eliceiri, “NIH Image to ImageJ: 25 years of image analysis,” Nature Methods, vol. 9, pp. 671–5, 2012.

[16] S. Culley, K. L. Tosheva, P M. Pereira and R. Henriques, “SRRF: Universal live-cell superresolution microscopy,” The International Journal of Biochemistry & Cell Biology, vol. 101, p. 74–79., 2018.

[17] T. Dertinger, R. Colyer, G. Iyer, S. Weiss and J. Enderlein, “Fast, background-free, 3D superresolution optical fluctuation imaging (SOFI),” Proceedings of the National Academy of Sciences, vol. 106, no. 52, p. 22287–22292, 2009.

[18] D. Bradley and G. Roth, “Adapting Thresholding Using the Integral Image,” Journal of Graphics Tools, vol. 12, no. 2, pp. 13–21, 2007.

[19] L. Zhu, W. Zhang, D. Elnatan and B. Huang, “Faster STORM using compressed sensing,” Nature Methods, vol. 9, pp. 721–3, 2012.

[20] E. Mukamel, H. Babcock and X. Zhuang, “Statistical deconvolution for superresolution fluorescence microscopy.,” Biophysical Journal, vol. 102, no. 10, pp. 2391–400, 2012.

