## Supplementary Information for "Sub-diffraction error mapping for localization microscopy images"

### Supplementary Note 1: Intensity based comparisons (SQUIRREL)

Identifying artefacts in a super-resolution reconstruction without ground truth information or any prior assumptions about the structure is a considerable challenge. The algorithms / methods used to produce the reconstructions may depend on such assumptions, a common one being local sparsity of the structure (as with Compressed Sensing [S1] and DeconSTORM [S2] used here). A significant first step towards this goal was the realisation by Culley et al [S3] that a limited amount of ground truth information for comparison could be extracted from the wide-field image. In their SQUIRREL algorithm [S3] the super-resolution reconstruction is down-sampled to produce an expected wide-field image by convolution with a resolution scaling function. How well the expected wide-field correlates with a reference wide-field image gives a measure of the fidelity of the reconstruction. The reference image can be acquired in a separate measurement or, in the case of localization microscopy, produced by summing all the camera frames of the super-resolution acquisition. The resolution scaling function assumes a linear relationship between the pixel magnitude in the super-resolution image and the photon intensities in the expected wide-field. It contains three necessary parameters that are optimised by minimising the difference between the expected and reference wide-field images. This approach, while useful for detecting other artefact, suffers from two major setbacks when trying to identify artificial sharpening.

Firstly, as the reconstruction is blurred up to the Point Spread Function (PSF) scale, almost all the fine structure information in the reconstruction is lost. Therefore, the results of different algorithms that produced different amounts of sharpening of small scale structure would appear almost identical after this transformation. Secondly, the assumption of a linear relationship between the intensity in the reconstruction and wide-field images can lead to incorrect ranking of reconstruction quality when the main artefact present is the artificial sharpening. The degree to which this relationship is true varies greatly between algorithms. An algorithm that is more linear but artefact prone may produce a lower error in reproducing the wide-field image than a less artefactual but less linear reconstruction. For example, we have found that an algorithm can often produce a worse error in SQUIRREL when HAWK pre-processing [S4] is used than when it is not, despite clearly producing a more accurate reconstruction. This problem is exacerbated by the necessary requirement to optimise the background and PSF in the Resolution Scaling Function (RSF). Tests on simulated data show that very small changes in the estimate of these parameters can reduce the measured error by more than the difference caused by a sharpened and authentic reconstruction. This can lead SQUIRREL to rank sharpened images above those that do not contain this artefact in certain circumstances (see Supplementary Fig. 1-2)

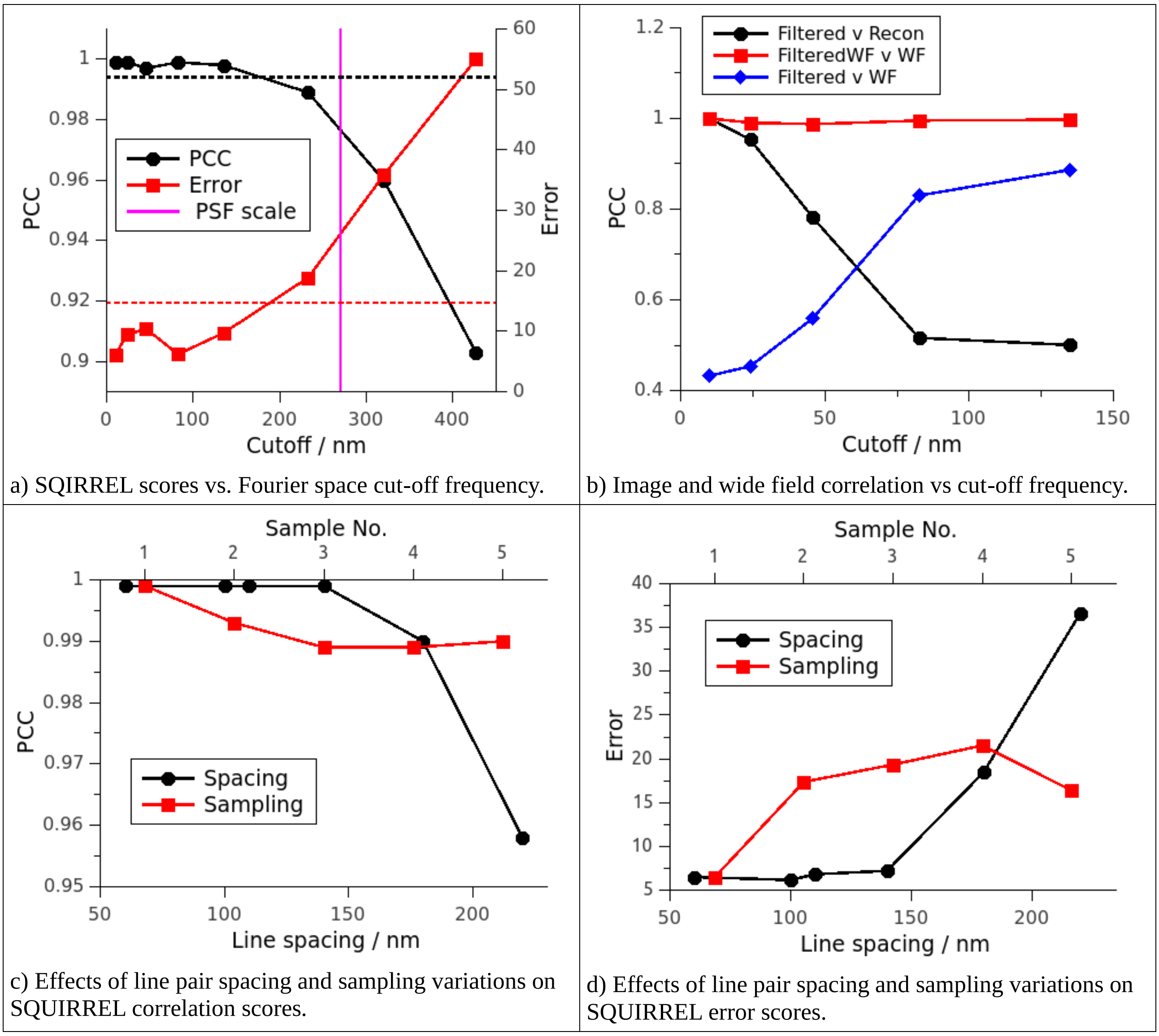


a) SQIRREL scores vs. Fourier space cut-off frequency.

b) Image and wide field correlation vs cut-off frequency.

c) Effects of line pair spacing and sampling variations on SQUIRREL correlation scores.

d) Effects of line pair spacing and sampling variations on SQUIRREL error scores.

**Supplementary Figure 1:** Insensitivity of SQUIRREL to errors in fine structure. Results of SQUIRREL analysis of a simulated low density dataset of a pair of lines separated by 100nm, where the sum of individual frames was used as the reference image. The fine structure of the reconstructions has been modified in two ways. a) & b) Fine structure was progressively removed by truncating the Fourier spectrum at larger ‘cut-off' frequencies (corresponding length on x axis) and taking the inverse Fourier transform as the input super-resolution image. a) The resolution scaled Pearson correlation coefficient (PCC) and resolution scaled error (Error) SQUIRREL scores show little variation with ‘cut-off' frequency until just below the point spread function (PSF) scale. Above this, variations are substantial, and SQUIRREL should reliably detect these artefacts. The scores produced by use of the ground truth (GT) as the test image are displayed by the dashed lines. The modified reconstructions score better than the ground truth significantly below the PSF scale despite containing substantial artefacts. b) A comparison of the correlation (PCC) of the input images at different scales. The filtered images show greatly reduced correlation with the unfiltered reconstruction as the ‘cut-off' frequency is increased (black). The removal of fine structure causes the reconstruction to more closely resemble the reference wide-field image (blue). Once these images are converted to wide-field scale images by applying a Gaussian blur and re-sampling (red), strong correlation is maintained and all sensitivity to artefacts is lost.

c) & d) Creation of artefacts by modifying the line spacing of the reconstruction. The spacing of the lines was increased or decreased by adding or removing columns of pixels to or from the centre of the two lines in the input reconstruction. SQUIRREL scores for correlation (c) and error (d) show almost no change with the size of the spacing until a little below the size of the PSF (black). For comparison the variation in scores for the unmodified reconstruction with the reference wide-field image produced from four other runs of the simulation (same structure and parameters) is shown in red. This variation is substantially larger than the variation due to line spacing below the PSF scale. This along with the ground truth comparison in (a) indicate that variations in sampling (emitter brightness and number of appearances) have a much larger effect on the image correlation than fine structural details i.e. artefacts.

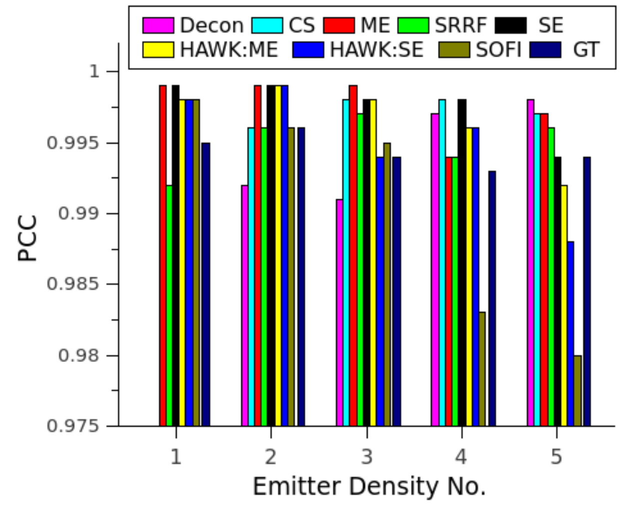


a) SQUIRREL correlation scores vs emitter density: 100nm line simulation.

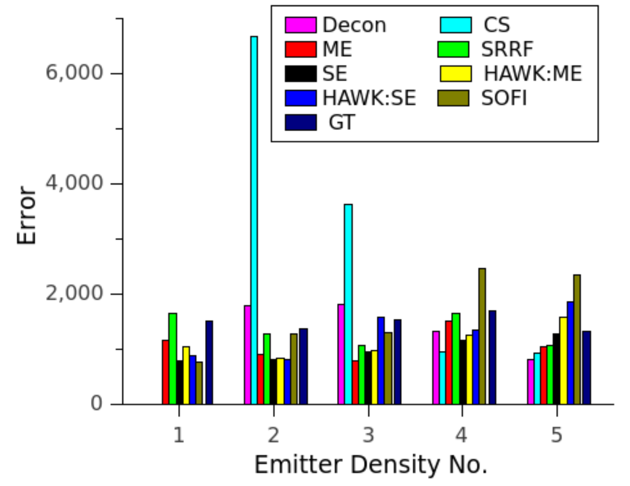


b) SQUIRREL error scores vs emitter density: 100nm line simulation.

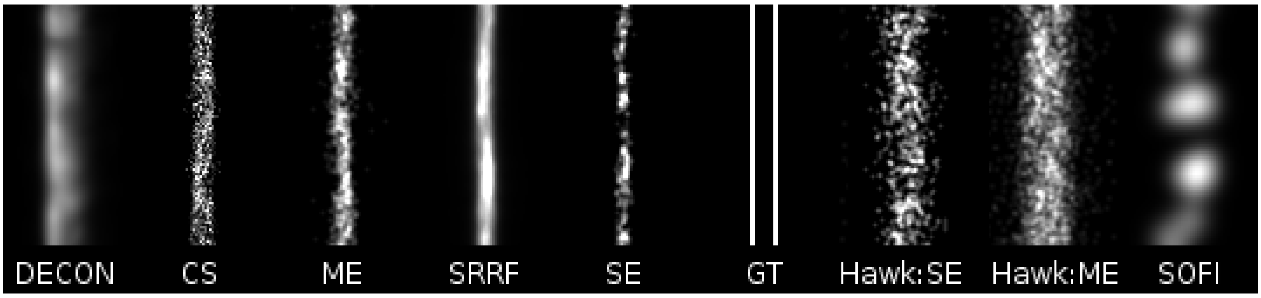


c) SQUIRREL ranking of reconstructions by PCC from best (left) to worst (right) for highest density simulation.

**Supplementary Figure 2:** Variation of the ability of SQUIRREL to detect artefacts produced by various high-density algorithms. The results of performing SQUIRREL analysis on simulated high emitter density data consisting of a pair of lines separated by 100nm using a number of specialist high-density algorithms. Simulations were produced at a range of emitter densities covering two orders of magnitude (No.1: low density - No.5: very high density). The resolution-scaled Pearson Correlation Coefficient scores (a) are very high for all algorithms [S1, S2, S4-S7], some scoring higher than the Ground Truth (due to the sampling/intensity effects described above). Only SOFI [S5] and to a lesser extent the two HAWK reconstructions show any significant fall off as emitter density increases despite being unable to resolve the structure. DeconSTORM, Compressed Sensing and Multi-Emitter fitting (ThunderSTORM) [S6] still give near perfect scores at the highest emitter density where they fail completely to resolve the structure. Similar trends are observed with the resolution scaled error score (b). The reconstructions at the highest emitter density are displayed in (c) ranked from best PCC score (left) to worst (right). Again, all but SOFI & HAWK score better than the Ground truth, despite obvious extreme sharpening/artefacts. HAWK and SOFI cannot resolve the spacing at this density but do (correctly) score worse than the GT and low-density results.

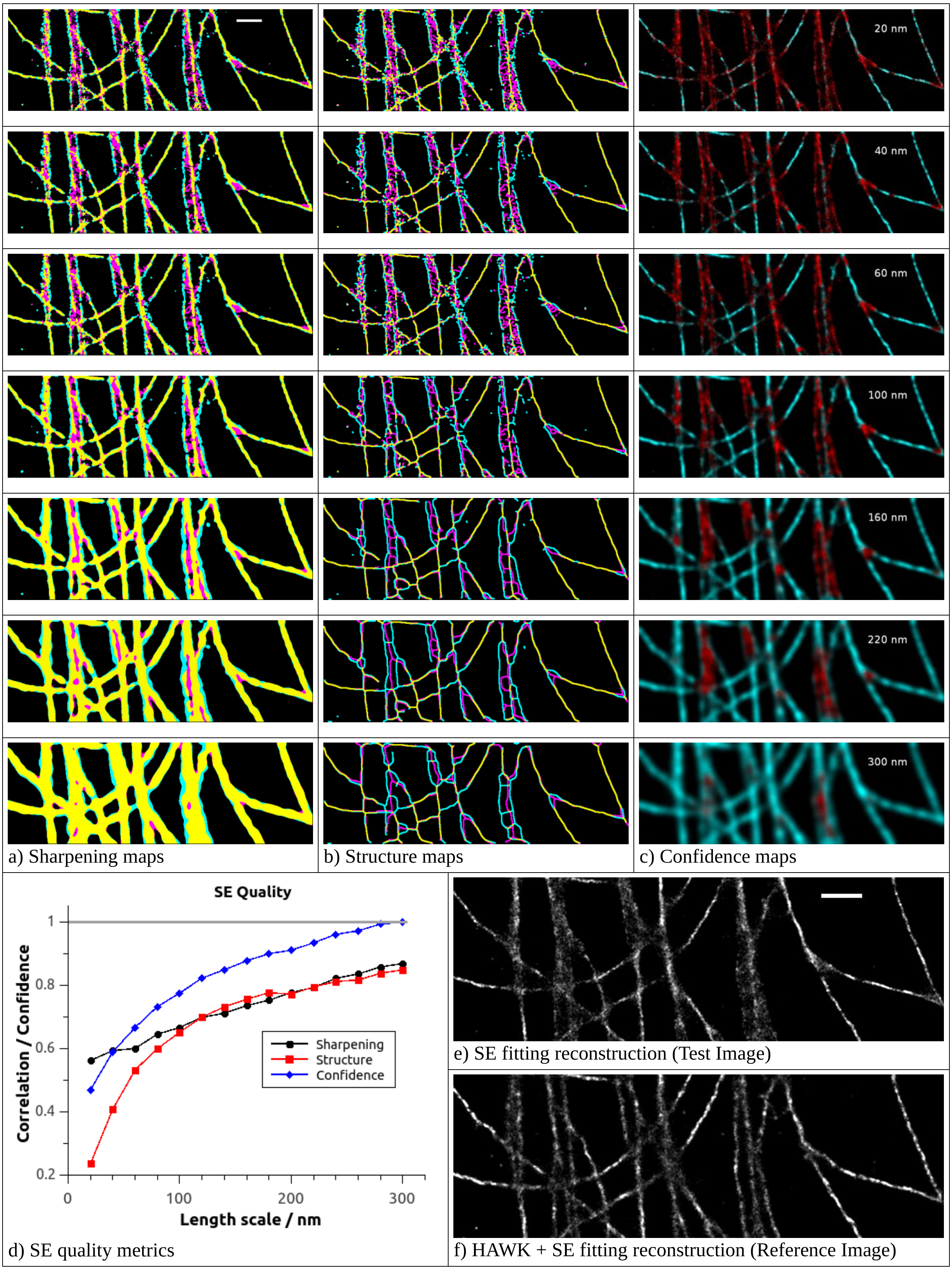


**Supplementary Figure 3:** HAWKMAN results for Single-Emitter (SE) fitting of microtubule data at various length-scales. (a-c) Sharpening, structure and confidence maps respectively for experimental microtubule data from the Localization Microscopy Challenge using the single emitter reconstruction as the test image and HAWK + SE as the reference. Results are shown for length scales 1, 2, 3, 5, 8, 11 & 15 pixels (20, 40, 60, 100, 160, 220 & 300 nm) from top to bottom. All three maps indicate substantial sharpening artefacts where microtubules are either closely spaced or crossing, as would be expected when the emitter density is too high. The first three scales (up to 60nm) show similar results, all indicating the degree of artificial sharpening in these regions is quite large, but limited elsewhere. At the 160nm scale (row 5) better agreement between test and reference is noted for both the sharpening and structure maps for some areas. The centre and left most microtubule bundles are now considered reliable parts of the reconstruction, as indicated in the corresponding confidence map. It should be noted the HAWKMAN can detect these artefacts even though the microtubules are not clearly resolvable in the HAWK reconstruction. The largest scale (300nm, bottom row) shows one area where the scale of sharpening in the test images is greater than the PSF (270nm).

The whole image correlation between test and reference sharpening and structure maps as a function of length-scale is shown in (d) (black & red respectively) along with the confidence metric used to colourise the confidence map (see methods) but calculated from the whole image (blue) rather than just the local correlation. These give an indication of the quality/reliability of the reconstruction as a whole at different scales. In this case the reconstruction is judged to be reliable only at around 300nm effective resolution. The unprocessed test & reference images are shown in e) & f) respectively, scale bars 1µm

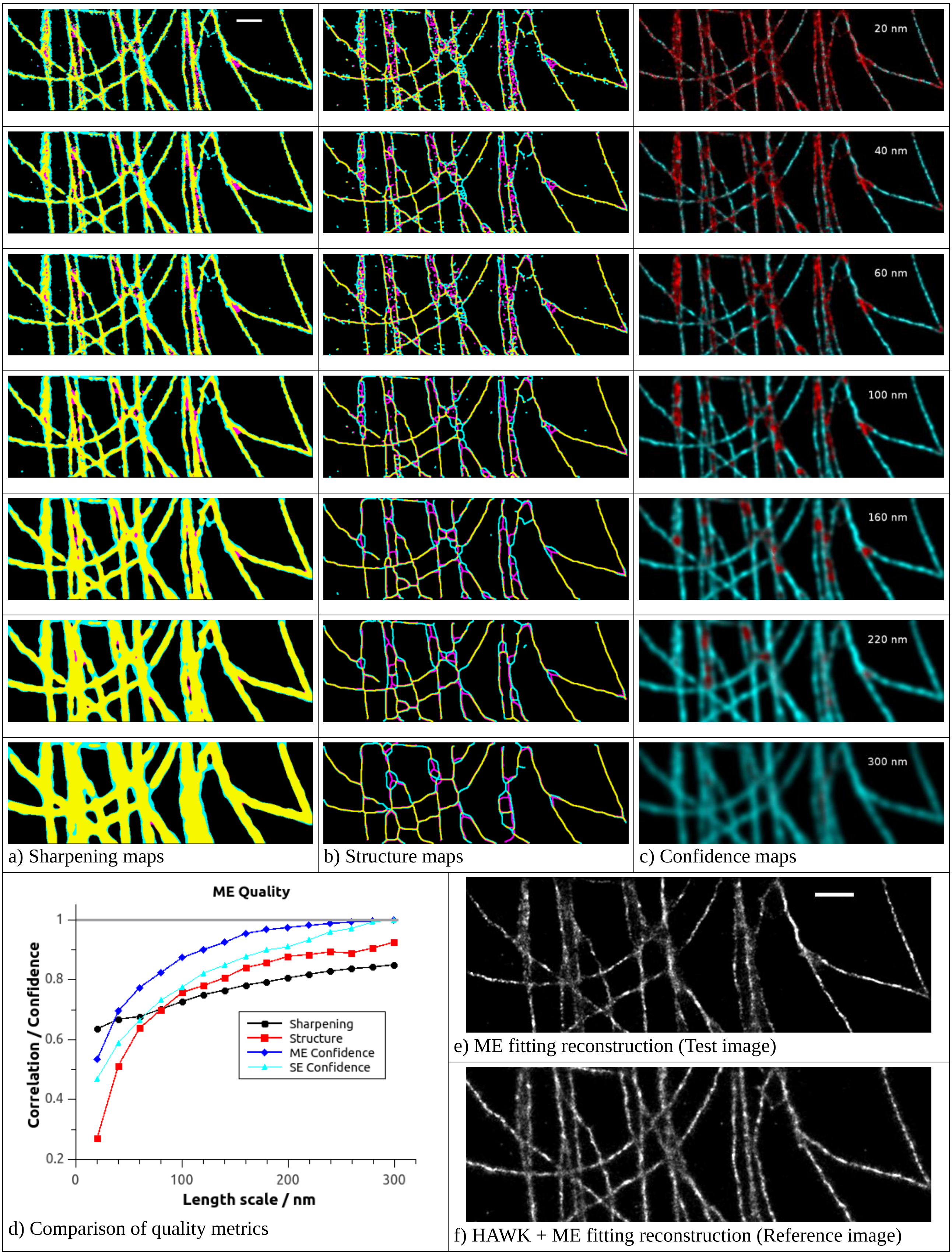


a) Sharpening maps b) Structure maps c) Confidence maps

d) Comparison of quality metrics

e) ME fitting reconstruction (Test image)

f) HAWK + ME fitting reconstruction (Reference image)

**Supplementary Figure 4:** HAWKMAN results for Multi-Emitter (ME) fitting of microtubule data at the same length scales as Single-Emitter (SE) fitting above. The same analysis as presented in Supplementary Figure 3 but with using the ME reconstruction as the test image and HAWK + ME as the reference. (a-c) The sharpening, structure and confidence maps at scales 20, 40, 60, 100, 160, 220 & 300nm (top – bottom). At scales up to 60nm (top 3 rows) the maps indicate that the ME reconstruction is only a modest improvement of the SE case. However, at length scales larger than this (100-220nm, rows 4-6) the improvement is substantial. The problematic areas of high structural density are indicated to be much more reliable. This can also be seen in the quality metrics (d). Here, the confidence metric from the SE result (cyan) which shows a significantly lower score compared to the ME result (blue) for intermediate length scales. Comparing the ME (e) and HAWK + ME (f) reconstructions (scale bar 1μm) with their SE counterparts in Supplementary Figure 3, shows it is these regions where the ME reconstruction is better resolved.

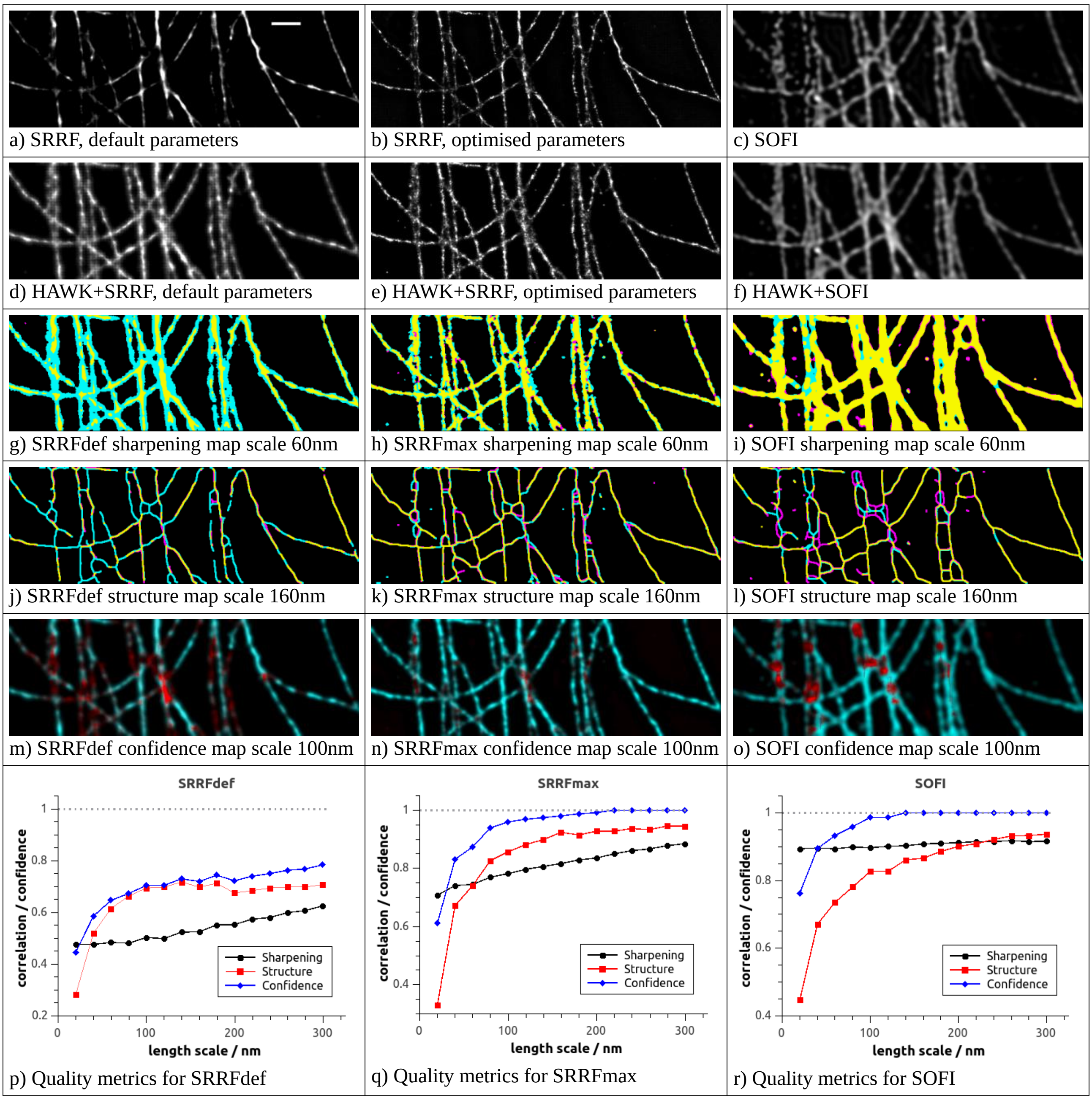


**Supplementary Figure 5:** HAWKMAN analysis on microtubule data from the Localization Microscopy Challenge using SOFI and SRRF [S7] as the test algorithm. (a-c) Reconstructions produced by the SRRF and SOFI algorithms without prior HAWK processing. a) The reconstruction produced by SRRF using the default parameters (SRRFdef – see methods), scale bar 1μm. b) SRRF reconstruction using previously optimised parameters (SRRFmax) including the Temporal Radiality Auto-Correlations (TRAC) option. Under appropriate conditions, these parameters should yield a higher resolution reconstruction [S7]. c) Reconstruction produced by 4th order SOFI analysis. (d-f) The corresponding reconstructions of each of these methods with prior HAWK processing for SRRFdef, SRRFmax and SOFI respectively. (a-c) were used as the test images and (d-f) as the reference images.

(g-o) Selected results for HAWKMAN analysis using these algorithms with and without HAWK for test and reference images respectively. For easier comparison the same length scales are displayed as for Figure 3 of the main text. (g-i) Sharpening maps for length scale of 3 pixels (60nm). (j-l) Structure maps for length scale 8 pixels (160nm). (m-o) Confidence maps for length scale 5 pixels (100nm). The corresponding quality metrics for the whole image for each algorithm are shown in (p-r). These consist of the correlation between test and reference sharpening maps (black), structure maps (red) and the corresponding confidence score calculated from these values (blue). The results for SRRF with default parameters taken together (g, j, m & p) show HAWKMAN is indicating the presence of substantial artefacts at all length scales. Comparing the sharpening and structure maps suggests these arise not just from collapse of adjacent structures to one, but also from the highly disjointed/punctate nature of the reconstructed microtubules and much sharper/finer structure in the test than in the reference image.

HAWKMAN results for SRRF with optimised parameters (h,k,n & q) suggest this reconstruction is of much higher quality. The test image (b) has much more continuous structure and individual microtubules are better resolved in the typical problem areas of high structural density. In this case the structure map (h) shows small differences in structure rather than just missing structure, and the sharpening map (k) shows only slight differences that are restricted to the problem areas. This suggests the precision estimated from the width of the structures would be much more accurate. From the confidence maps and scores (n & q) HAWKMAN is indicating this reconstruction is fairly accurate at the 100nm scale.

Results for the SOFI reconstruction (i,l,o & r) show very little sharpening (i) and only slight structural differences. This suggests the artefacts consist of disjointed rather than collapsed structure, again as a consequence of the highly non-linear relationship with intensity. The whole image confidence score suggests an accurate reconstruction down to the 100nm scale.

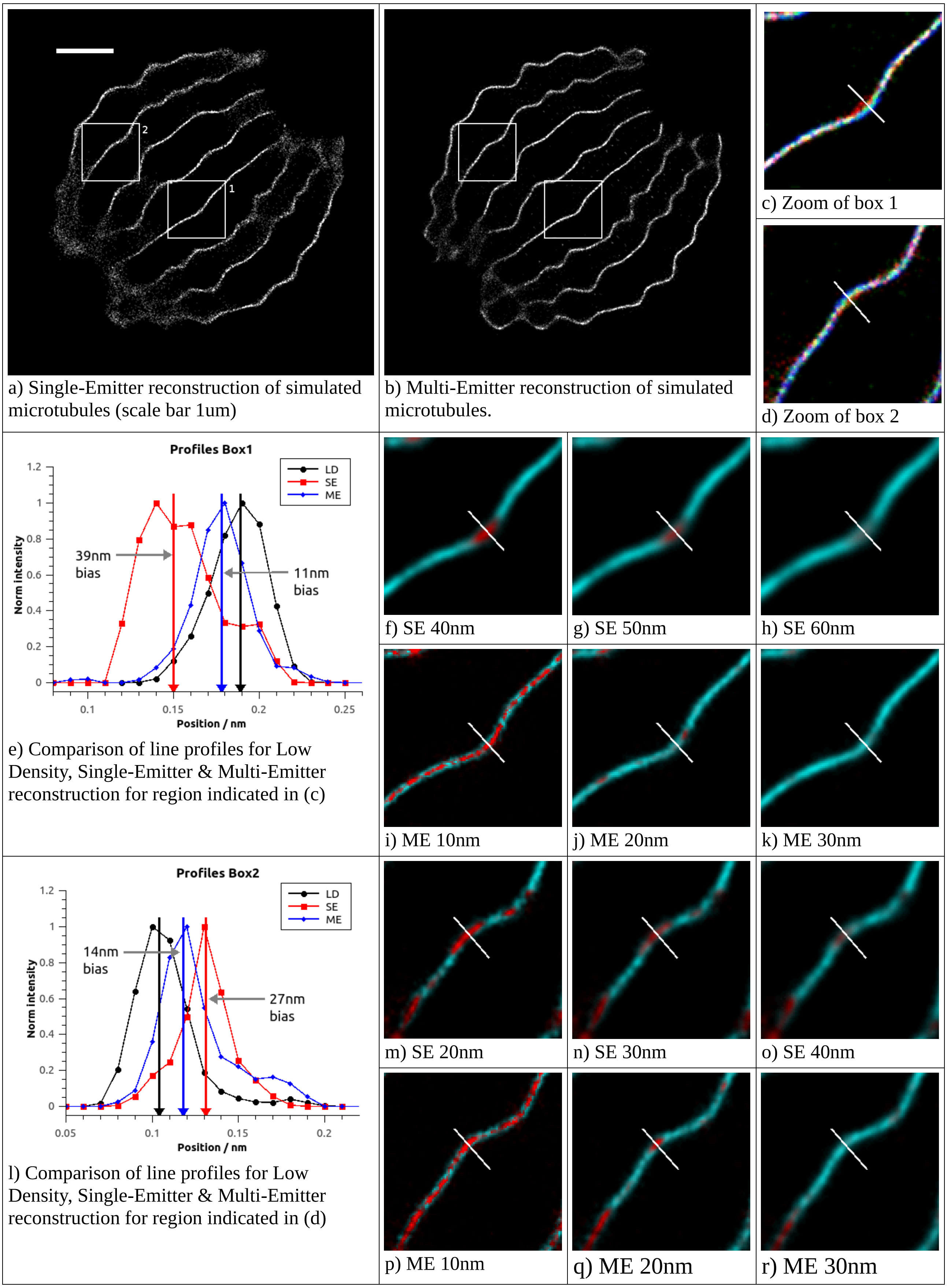


**Supplementary Figure 6:** Test of the ability of HAWKMAN to accurately determine the scale of small artefacts in simulated high-density microtubule data from the Localization Microscopy Challenge [S8]. (a, b) ThunderSTORM reconstructions of the high emitter density data using single-emitter (a) and multi-emitter (b) fitting, scale bar 1µm. Both reconstructions show obvious artefacts at the extremities where the structural density is highest. The boxed regions highlight areas of the reconstruction where the microtubules are well separated and show no obvious sharpening effects. Comparison between the two reconstructions, however, shows that in the Single-Emitter reconstruction the microtubules are significantly straighter than in the Multi-Emitter counterpart. An overlay of the two results with the reconstruction from the Low Density simulation is shown in c) & d) for boxes 1 & 2 respectively (SE-red, ME-green, LD-blue). These indicate a quite significant bias for the SE result compared to the LD reconstruction. The ME result is much closer to the LD but still contains some artefacts. These regions were selected because they display relatively small-scale artefacts. Other regions of the reconstructions display much larger scale differences, such as the more curved lowermost microtubule.

A selection of HAWKMAN results for the region in box 1 are shown in (e-k) for SE and ME reconstructions. The scale of the bias was estimated from line profiles measured perpendicular to the microtubule (indicated by line in (c), length 300nm). The measured profiles are displayed in (e). The vertical arrows indicate the central position of a Gaussian fit to each profile. The difference between these positions for the high and low density data quantified the bias indicated on the graph. HAWKMAN confidence maps are shown at selected scales for the SE (f-h) and ME (i-k) reconstructions. The SE maps (f-h) suggest a strong likelihood of artefacts at the 40nm scale, a moderate likelihood at the 50nm scale, but no artefacts at the 60nm scale or larger. This compares well to the measured scale of 39nm. For ME fitting, the maps (i-k) suggest artefacts of 10nm scale but none of 20nm scale, comparing well to the measured 11nm (from fitted line-scans) for the ME reconstruction.

Similar results are observed for the region in Box 2 (l-r). Here the SE maps (m-o) suggest artefacts of scale 30nm and possibly 40 nm compared to a measured value of 27nm. The ME maps (p-r) indicate strong likelihood of artefacts at 10nm, moderate likelihood at 20nm but none at 30nm. Again, this compares favourably with the measured value of 14nm.

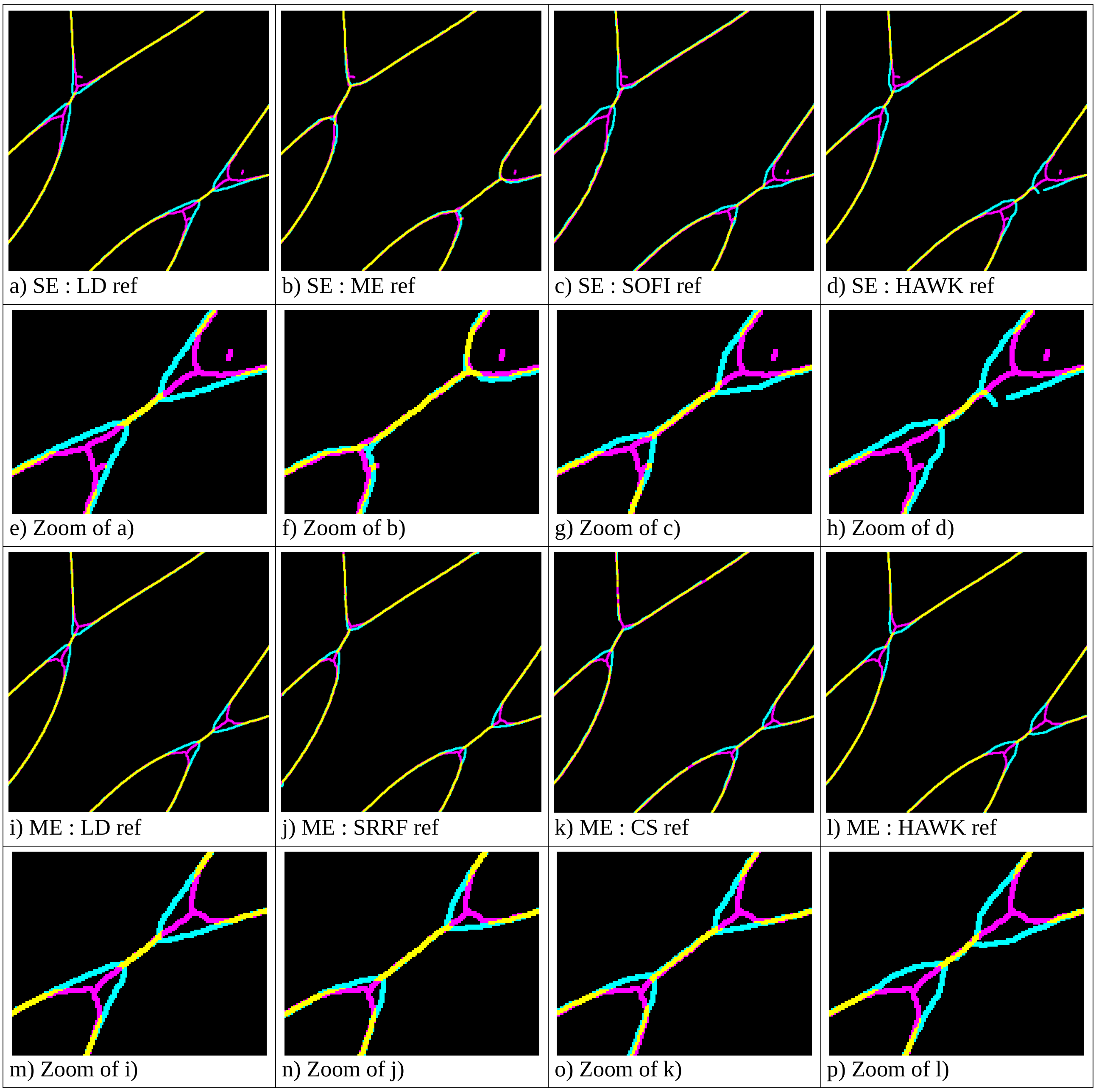


**Supplementary Figure 7:** Performance comparison when using different algorithms for the reference image. Here the ability of HAWKMAN to identify sharpening artefacts is demonstrated on simulated microtubule-like data from the Single Molecule Localization Microscopy Challenge [S8]. (a-d) show the skeletonised output of tests on the Single-Emitter (SE) fitting reconstruction (using ThunderSTORM [S6]. Magenta = Test image only, cyan = Reference only, yellow = agreement between both) using various other algorithms for the reference image. Enlarged regions of (a-d) are shown in (e-h) respectively. Results are displayed for length scale = 8 SR pixels (80nm). Dilation (ImageJ) has been applied to the images for clarity of display.

(a, e) Use of the low-density SE reconstruction of the same structures for the reference image provides an accurate, artefact free image for comparison. Any deviation between this image and the test image highlights sharpening artefacts present in the latter, and represents the optimum reference image for artefact detection. Severe disagreement in the structure (sharpening) between the test (SE) and reference (low-density) reconstructions is present either side of where the microtubules cross, indicating strong sharpening artefacts. (b, f) When the Multi-Emitter (ME) fitting reconstruction is used as the reference, the skeletonised images are almost identical and no significant artefacts are revealed. This is because both images are equally sharpened at this scale. (c, g) SOFI performs significantly better as the reference despite being (in principle) a lower resolution reconstruction. It detects some of the sharpening present, but not the regions closer to the crossing point. (d, h) Using the HAWK pre-processed reconstruction for the reference gives results very similar to the low-density case, revealing all of the sharpening artefacts present in the test image. Using either Compressed Sensing or SRRF [S7] for the reference (not shown) gives results intermediate between ME fitting and SOFI.

(i-p) Same as above but with ME fitting used for the test image. Again, using HAWK:ME for the reference image greatly outperforms other algorithms (CS and SRRF) and gives results very similar to using the low-density reconstruction for the reference, indicating detection of all artefacts.

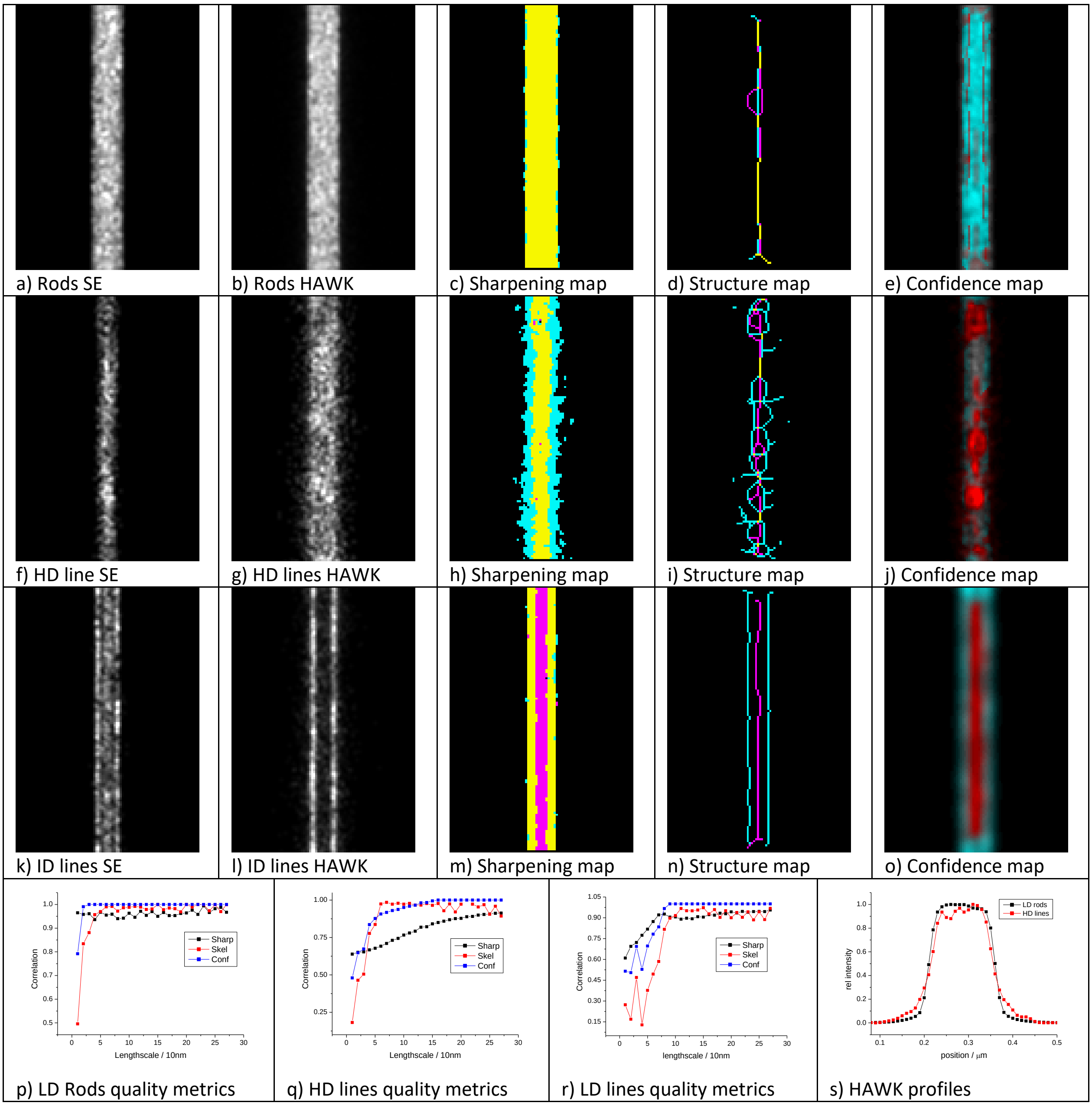


**Supplementary Figure 8:** HAWKMAN can also be used to assess the quality of the HAWK reference image. Here three different structures are simulated at different emitter densities to demonstrate the different types of artefact detected with HAWKMAN. The top row (a-e) show results for a 150nm wide planar structure with an even labelling density across its width, simulated at a low emitter density. Consequently, the unprocessed (a) and HAWK pre-processed (b) Single Emitter reconstructions appear similar. HAWKMAN results (representative length scales, c-e) highlight these structural similarities. Both the sharpening map (c,10nm length scale) and the structure map (d. 30nm length scale), show close agreement between test (a) and reference (b) images. The confidence map (c, 20nm length scale) shows a high confidence score throughout the structure.

The second row shows results for a line pair structure with a separation of 100nm at very high emitter density, sufficient for the spacing to be unresolved by SE fitting even when HAWK is used. The test image (f) is extremely sharpened, whereas the HAWK+SE reference (g) shows substantial loss of precision. The density was tuned such that the reduced precision results in a HAWK+SE reconstruction that closely resembles the 150nm planar simulation (b). Visual inspection alone would not be able to identify which of these reconstructions (b,g) was a more authentic representation of the underlying structure. HAWKMAN dramatically highlights the difference (h-j). The sharpening map (h, 10nm length scale) show the precision loss as large areas of reference only structure (cyan) bordering areas of common agreement (yellow). The structure map (i, 30nm length scale) shows a much more branched structure for the reference (cyan) than the test image (magenta) which we have observed to frequently be associated with unresolved structure in the refence. The 20nm length scale confidence map (j) therefore shows the presence of substantial artefacts in the test image at this scale. While the HAWK+SE reference does not contain the extreme sharpening bias of the test image, the HAWKMAN sharpening & structure maps suggest that the precision of the HAWK+SE reference is insufficient to guarantee there is no unresolved fine structure of this scale (10-30nm). There would therefore likely to be significant improvement from using HAWK with Multi-Emitter fitting.

This contrast strongly with the case of the same structure simulated at an intermediate emitter density (3$^{rd}$ row) where the SE test image is sharpened (k) but the gap is clearly resolved in the HAWK+SE reference (l). The sharpening map (m, 30nm scale) indicates no significant loss of precision (cyan boarders) in the reference just substantial false structure(magenta) in the test reconstruction. Similarly, the structure map (n, 40nm scale) clearly shows structure that is resolved in the reference that is unresolved in the test. The confidence map (o, 70nm scale) correctly indicates where the reconstructed structures are authentic. These results suggest HAWK+SE fitting is adequate and only modest improvements would be expected by using HAWK with ME fitting.

HAWKMAN quality metrics for the above simulations over the full range of length scales are shown in (p-r). For the low-density simulation (a-e) these metrics indicate a highly authentic reconstructions (p) at all but the 10nm length scale (limit of localization precision). For the very high-density simulation (f-j, q) the sharpening maps show poor correlation (black) even for length scales significantly larger than the 100nm line separation, whereas the structure maps correlate well for length scales greater than 50nm. We have frequently observed this discrepancy when the emitter density is sufficiently extreme to significantly reduce the localization precision in the HAWK reference. This further suggests the true structure may contain elements at smaller length scales that on not resolved in reference image (which in this case it does). For the intermediate density simulation (k-o,r) then correlation of the sharpening maps (black) and structure maps (red ) roll off at a similar length scale, suggesting the HAWK reference is probably authentic (which it is). A comparison of the intensity line profiles (s) of the HAWK+SE reference reconstructions for the low-density planer structure (black) and the high-density line structures (red) shows just how similar these structures appear in the reconstructions despite their actual substantial differences in structure.

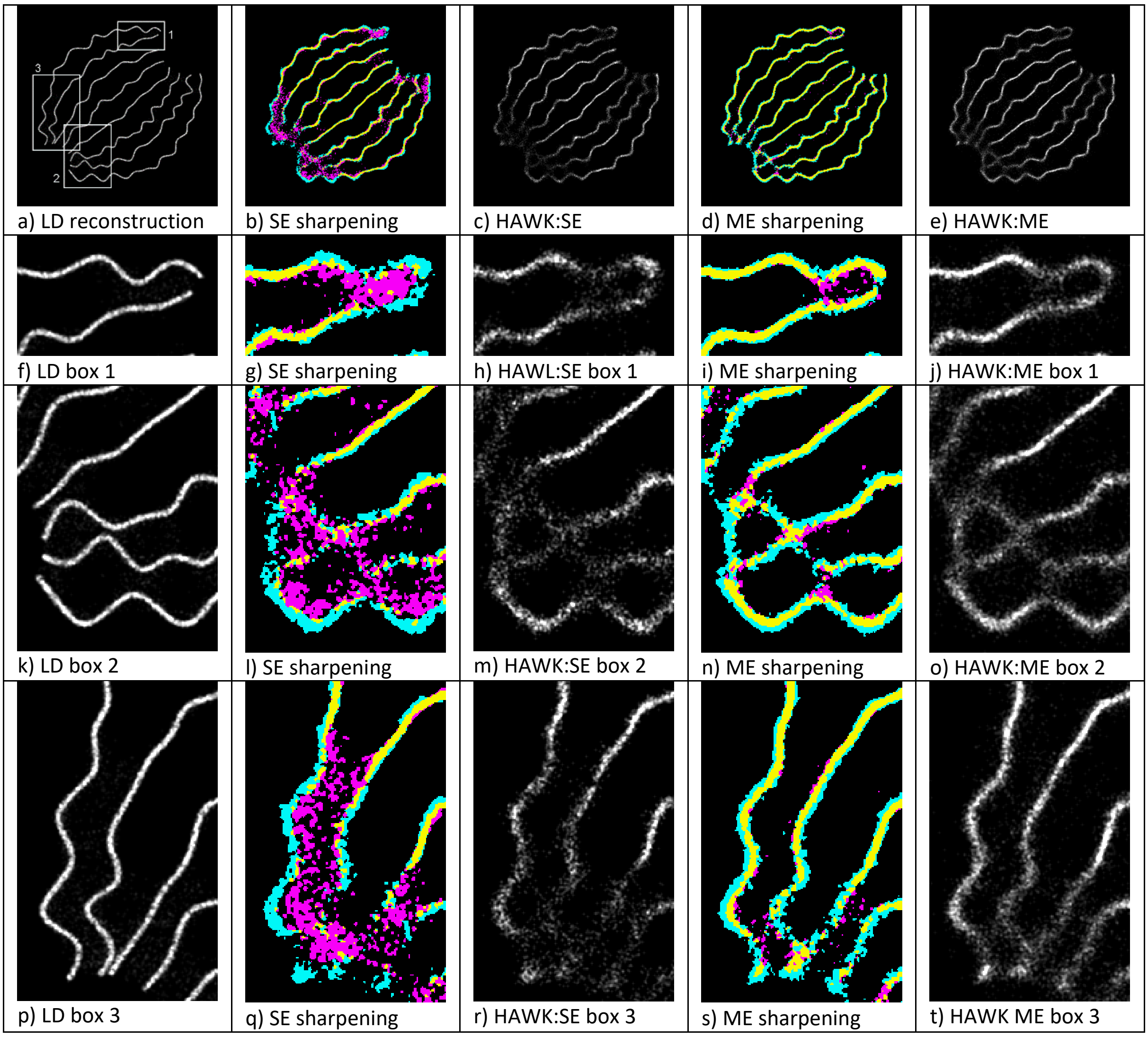


**Supplementary Figure 9:** Evaluation of HAWK precision on alternative simulations from the Localization Microscopy challenge [S8]. The ability of HAWKMAN to assess the precision of the HAWK reference can be tested by comparing its predictions with a low-density reconstruction. (a) Single Emitter reconstruction of low-density simulation. (b) HAWKMAN sharpening map (20nm length scale) for SE fitting of a high-density simulation showing substantial structural differences between test and reference images. These are particularly prevalent at the edges of the image where the fibres converge with the cyan and magenta areas highlighting missing and false structure in the test image. These areas show a relatively lower localization precision in the HAWK+SE reconstruction (c). The comparable sharpening map for Multi Emitter fitting (d) shows much improvement in these areas and the corresponding HAWK+ME reference image (e) a much more consistent level of precision throughout the reconstruction.

Close up views of all the images for the boxed sections in (a) are shown in (f-j) for box 1, (k-o) for box 2 and (p-t) for box 3. The sharpening map for box 1 (g) show substantial sharpening for the SE test image (magenta) where the fibres converge on the righthand side. The HAWK+SE reference only structure (cyan) shows much greater accuracy compared to the low-density reconstruction (f). These cyan areas also show where the emitter density is likely sufficiently high that the precision of the HAWK reference will be noticeably reduced. This is confirmed by comparing the high density HAWK+SE reconstruction (h) to the low density image (f), which shows increased scatter in the areas marked by cyan in the sharpening map, but much more similar precision where there is some common structure between the high density test and references agree (yellow, left-hand side). The sharpening map for ME fitting (i) indicates moderate sharpening (bias, magenta) in the test image and only limited precision loss in the HAWK+ME reference in one specific area. This is replicated in the reference image (j) that show a much more consistent level of scatter throughout the reconstruction and compares more favourable to the low-density reconstruction.

The SE reconstructions for box 2 and box 3 show similar results (l-m & q-r) as above with substantial sharpening of the test (SE only) reconstruction (magenta) and the cyan only areas predicting where the precision in the reference (HAWK+SE) is significantly reduced. For the ME fitting results (n-o,s-t) more subtle sharpening of the test (ME only) image is observed, both by the magenta images and where the common structure(yellow) is not located centrally in a cyan area, indicating bias. Where this is the case, a modest reduction in precision is observed in the HAWK+ME reconstruction (t), whereas the cyan (reference only) areas or the sharpening map are associated with a more significant increase is scatter in the HAWK+ME reference.

These results combined suggest that not only is the HAWK pre-processed reconstruction always superior to when HAWK is not used for both single and multi-emitter fitting (as expected). Additionally, they also suggest that HAWKMAN can give an indication of where in a HAWK pre-processed reconstruction the fidelity (precision and accuracy) approaches that which would be obtained in a low-density acquisition and where it is significantly below this.

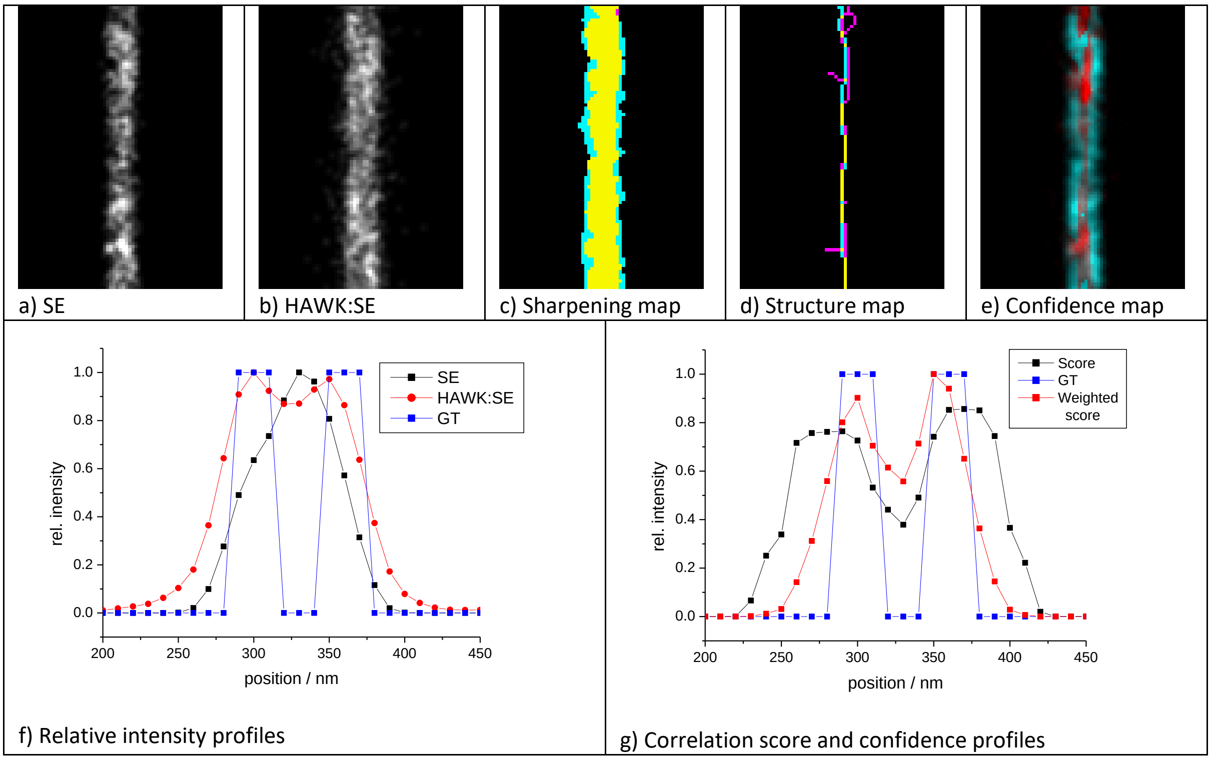


**Supplementary Figure 10:** HAWKMAN analysis of synthetic data simulating Z1Z2 antibody labelling of muscle sarcomeres, allowing for spatial variations due to antibody size and structural inhomogeneity in the z axis. These effects contribute to a line width (20nm) significant compared to the line spacing). Simulations are at a density to cause significant but not extreme sharpening. This combined with a high background and low signal to noise (as in the experimental data) lead to the spacing being unresolved in the Single Emitter reconstruction (a). The high background and low signal mean the spacing is still not properly resolvable when HAWK is used (b). but does indicate a slightly wider structure. This is reflected in the HAWKMAN sharpening map (c, 20nm length scale) where the significant cyan areas bordering the structure indicate the possibility of a sharpening artefact. The branched nature of the structure map (d), also at 20nm length scale, suggests the possibility of unresolved structure. The confidence map (e, 20nm) also indicates the structure may actually be split into a doublet of lines, with low confidence score for the central part (red).

These inferences are further supported by the intensity profiles across the images (g). Here the standard SE reconstruction (black) cannot resolve the spacing. The HAWK:SE profile (red) only suggest the possibility that the structure is in fact made of two lines but gives a much better indication of the overall size when compared to the Ground Truth (blue). A stronger indication that there may be an unresolved spacing is seen by how the HAWKMAN metrics vary across the images (g). The combined correlation score (black) indicates a much lower confidence in the central part of the reconstruction where there is no real structure. When weighted by the intensity (red) this gives a clear indication of the presence of a split structure. In this specific case, the spacing indicated by this metric is in close agreement with Ground truth (blue) but this may be a unique feature of parallel line structures. However, for other structures HAWKMAN should still give an indication of when structure might be missing from both the test and reference images due to excessive artificial sharpening and precision loss.

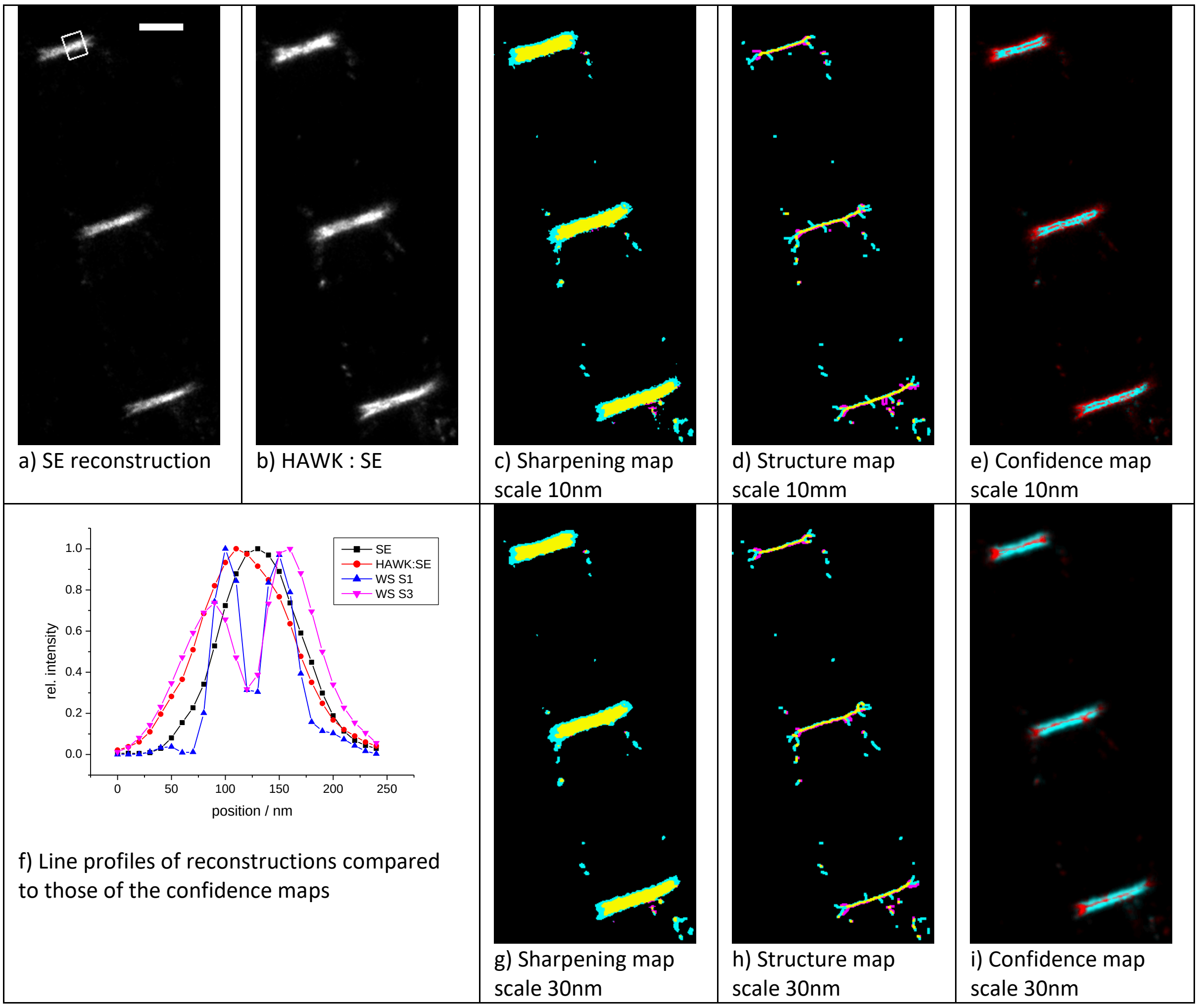


**Supplementary Figure 11:** HAWKMAN analysis of experimental data of Z1Z2 antibody labelling of muscle sarcomeres. Data were collected at sufficient emitter density to produce significant artificial sharpening when using Single Emitter fitting. This combined with other possible effects (structural variation, antibody labelling, signal to noise and drift) mean the expected line separation is not resolved in either the unprocessed (a) or HAWK (b) reconstructions. HAWKMAN analysis (c-e) at the 10nm scale suggest the possibility of missing structure. The Sharpening map (c) indicates some loss of precision in the reference image (cyan areas). The highly branched structure map (d) is indicative of partially resolvable structural differences in the two images. The confidence map (e) highlights the lack of complete correlation between the structures in the input images, suggesting some sharpening artefacts in the test image and the possible presence of unresolved structure in the HAWK reference.

Comparison of the line profiles across the boxed region in (a) are displayed on the graph in (f). The unprocessed test (black) and HAWK pre-processed reference (red) intensity profiles show some moderate differences, indicating sharpening artefacts in the test and possible loss of precision in the reference. The intensity weighted correlation score (see above simulation example in Supplementary Fig. 10 for definition) show a greatly reduced confidence in the central part of the structure for both the 10nm (blue) and 30nm (magenta) length scales. These two profiles both show a bimodal distribution similar to the simulated data above with a peak spacing of ca 50 nm (10nm length scale) and 70nm (30nm length scale). Although these distances compare favourably with the expected separation (60-70nm), they differ significantly from each other over a short range of length scales. This combined with the limited information contained in the input images along with the arguments given for the analysis of simulated data above mean that this should not be interpreted as a reliable measure of the underlying structure, but merely that underlying structure of some form may well exist. (g-i) show the 30nm length scale results for the sharpening, structure and confidence maps respectively. The confidence map shows how HAWKMAN consistently rejects all these structures as unreliable. Scale bar 500nm.

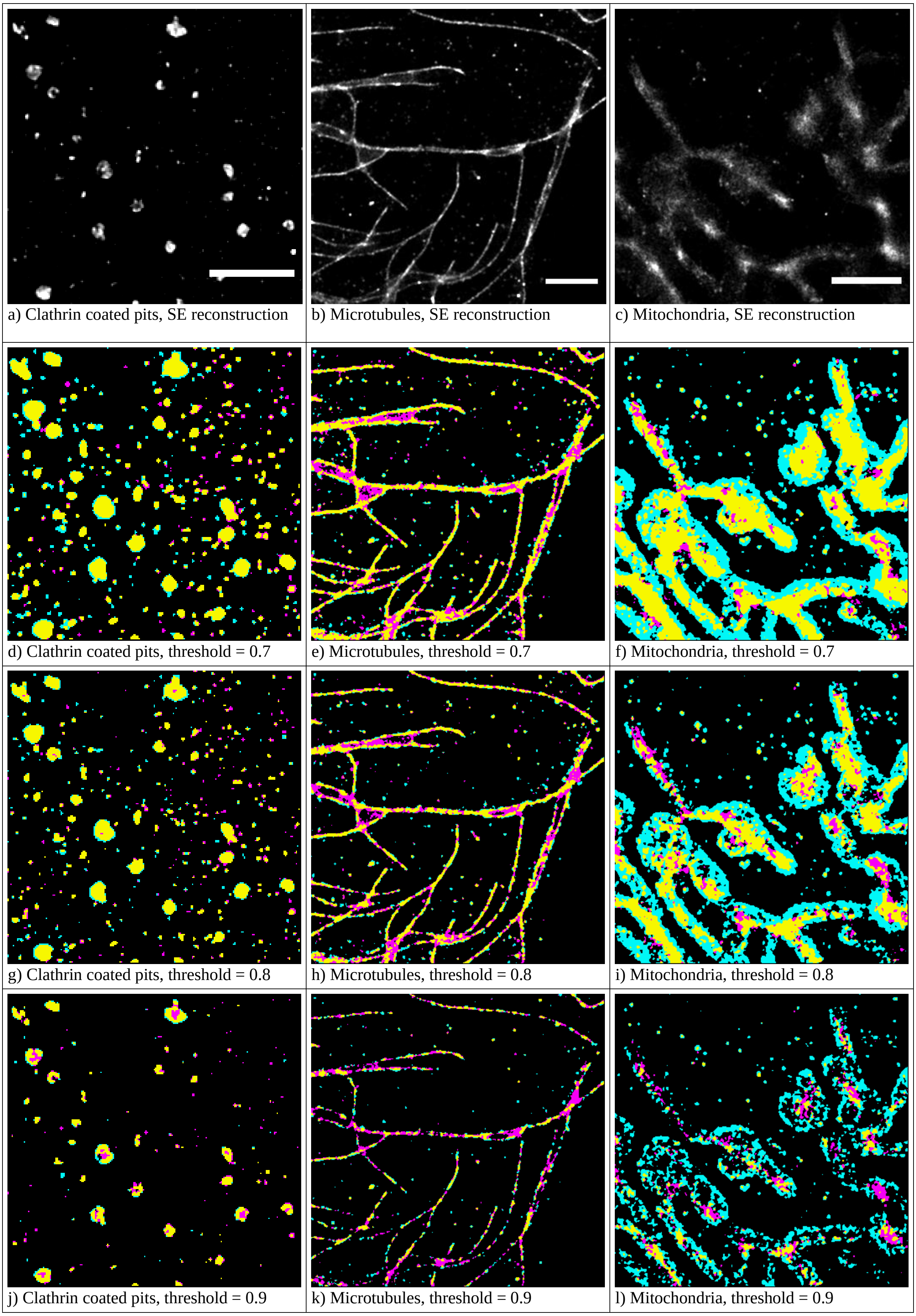


**Supplementary Figure 12:** The effects of local threshold on ability of HAWKMAN to detect artefacts. HAWKMAN is generally not very sensitive to the choice of threshold parameters, but the optimum value can be anticipated from the nature of the structures. (a-c) Single-Emitter (SE) reconstructions for clathrin-coated pit, microtubule and mitochondria data respectively. (d-f) HAWKMAN sharpening maps (length scales 32nm, 32nm & 96nm respectively) produced using the default threshold coefficient of 0.7. The depth of the 'hole' in the clathrin coated pits (d) is too shallow to be detected using this value as it is intended to detect features with an intensity contrast of approximately a factor of 2. The contrast between centre and edge is less than this for the pits in the image. For microtubules (e) and mitochondria (f) that have greater contrast, HAWKMAN reliably detects artefacts with this default value. Very little difference is observed with a higher threshold coefficient of 0.8 (g-i). With a coefficient of 0.9 (j-l) HAWKMAN easily detects the artefactual infilling of the shallow hole in the clathrin coated pits (j) but for the microtubule and mitochondria data (k & l), this high value leads to some missing structure in the HAWKMAN maps, but still identifies similar areas as containing artefacts as with the default value. Scale bars are 2µm

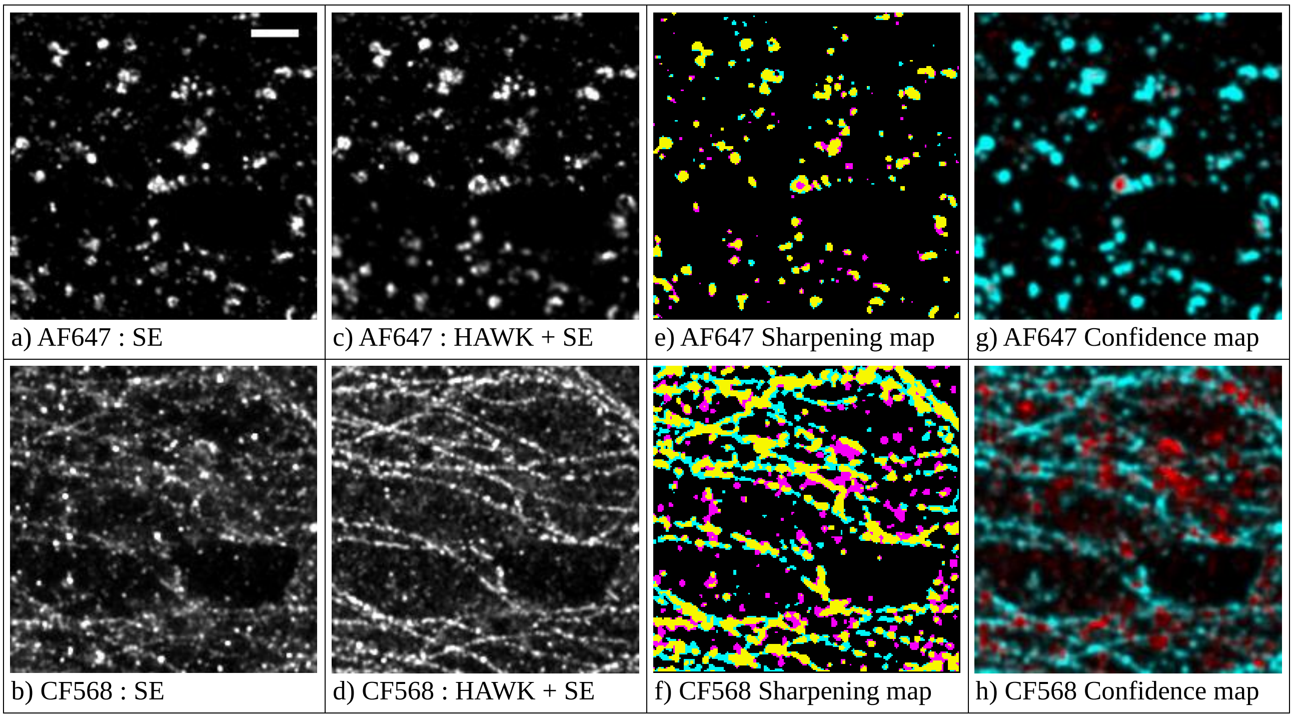


**Supplementary Figure 13:** HAWKMAN is sensitive to artefacts likely to be encountered in two colour imaging. (a, b) Single-emitter reconstructions for clathrin-coated pits stained with Alexa Fluor 647 (a, lower emitter density) and tubulin stained with CF568 (b, higher emitter density) were compared using HAWKMAN with their HAWK pre-processed counterparts (c, d) to evaluate image fidelity. HAWKMAN sharpening maps (e, f) were generated using structure-appropriate threshold coefficients of 0.9 for clathrin and the default value of 0.7 for tubulin. Confidence maps (g, h) were produced using a correlation measure of the blurred sharpening maps and the blurred structure maps (not displayed), the latter of which were used a threshold coefficient of 0.95 in the clathrin-AF647 case (a, c, e, g) and a default value of 0.85 in the tubulin-CF568 case (b, d, f, h). Due to high background, the global offset coefficient was also raised from its default value of 0.04 to 0.1 in the tubulin-CF568 case. Very little sharpening is present in the low-density AF647-clathrin case (a, c, d, g), as evidenced by the prevalence of cyan colouring in (g) and yellow dominating all non-background signal in (e), except for a single clathrin-coated pit whose 'pit' is unresolvable in the non-HAWK case (a) that is detected by both HAWKMAN maps (e, g). This demonstrates the value of HAWKMAN even in cases where a low density of emissions would often be assumed to result in little sharpening. In the high-density case, the sharpening map (f) shows the source of the severely punctate nature of the test image (b) to be a conflation of emissions towards their mutual centre both between (f, magenta) and along (f, cyan) the microtubules, with magenta colouring suggesting an introduction of false structure between proximal fibres, and cyan suggesting an absence of data due to emitter mis-localization. All maps are rendered for analysis at a 96nm length scale (3 pixels at 5x magnified reconstruction pixel size of data taken on a 160nm camera). Scale bar is 1 µm.

### Supplementary References:


[S1] Zhu, L., Zhang, W., Elnatan, D., and Huang, B. Faster STORM using compressed sensing. *Nature Methods* **9**, 721–723 (2012).

[S2] Mukamel, E. A., Babcock, H., and Zhuang, X. Statistical Deconvolution for Superresolution Fluorescence Microscopy. *Biophys J.* **102**(10) 2391-2400 (2012).

[S3] Culley, S., Albrecht, D., Jacobs, C., Pereira, P. M., Leterrier, C., Mercer, J. and Henriques, R. NanoJ-SQUIRREL: quantitative mapping and minimisation of super-resolution optical imaging artefacts. *Nature Methods* **15**(4) 263-266 (2018).

[S4] R. J. Marsh, K. Pfisterer, P. Bennett, L. M. Hirvonen, M. Gautel, G. E. Jones and S. Cox, "Artifact-free high-density localization microscopy analysis," *Nature Methods*, p. 689–692, 2018.

[S5] T. Dertinger, R. Colyer, G. Iyer, S. Weiss and J. Enderlein, "Fast, background-free, 3D super-resolution optical fluctuation imaging (SOFI)," *Proceedings of the National Academy of Sciences*, vol. 106, no. 52, p. 22287–22292, 2009.

[S6] M. Ovesný, P. Křížek, J. Borkovec, Z. Švindrych and G. M. Hagen, "ThunderSTORM: a comprehensive ImageJ plug-in for PALM and STORM data analysis and super-resolution imaging," *Bioinformatics*, p. 2389–90, 2014.

[S7] Gustafsson, N. et al. Fast live-cell conventional fluorophore nanoscopy with ImageJ through super-resolution radial fluctuations. *Nature Communications* **7**, 12471 (2016).

[S8] D. Sage, H. Kirshner, T. Pengo, N. Stuurman, J. Min, S. Manley and M. Unser, "Quantitative evaluation of software packages for single-molecule localization microscopy," *Nature Methods*, vol. 12, no. 8, pp. 717-29, 2015.

### Supplementary Videos

**Supplementary Video 1:**

Full results of HAWKMAN analysis of the Localization Microscopy Challenge microtubule data using Single-Emitter fitting in ThunderSTORM. Top to bottom show the sharpening map, the structure map and the confidence map respectively. Each frame corresponds to a 20nm increase in the length scale used, ranging from 20nm (frame 1) to 300nm (frame15). Colours and scale are as indicated in Fig.1 of the main text. For clarity of display one level of dilation has been applied to the structure maps.

**Supplementary Video 2:**

Full results of HAWKMAN analysis of the Localization Microscopy Challenge microtubule data using Multi-Emitter fitting in ThunderSTORM. Colours, maps, frames and scale are as Supplementary Video 1. For clarity of display one level of dilation has been applied to the structure maps.

**Supplementary Video 3:**

Full results of HAWKMAN analysis of the Localization Microscopy Challenge microtubule data using the SRRF algorithm with default parameters (see methods). Colours, maps, frames and scale are as Supplementary Video 1. For clarity of display one level of dilation has been applied to the structure maps.

**Supplementary Video 4:**

Full results of HAWKMAN analysis of the Localization Microscopy Challenge microtubule data using the SRRF algorithm with optimised parameters (see methods). Colours, maps, frames and scale are as Supplementary Video 1. For clarity of display one level of dilation has been applied to the structure maps.

**Supplementary Video 5:**

Full results of HAWKMAN analysis of the Localization Microscopy Challenge microtubule data using 4th order SOFI. Colours, maps, frames and scale are as Supplementary Video 1. For clarity of display one level of dilation has been applied to the structure maps.
