## Supplementary material for "Sub-diffraction error mapping for localization microscopy images": Plugin: ReadMe.txt

****************************************************************
COX GROUP PLUGIN - ANALYSIS SOFTWARE FOR LOCALISATION MICROSCOPY
****************************************************************

A Plugin for ImageJ containing various tools for the analysis of Localisation microscopy data and images
Package includes :

	HAWK v 1.3 	- A tool for pre-processing Localisation microscopy image sequences prior to localisation
			Prevents excess emitter density from producing artificial sharpening artifacts in the reconstruction.

	
	HAWKMAN v 1.0	- A tool for assessing the accuracy and identifying artefacts in Localisation Microscopy reconstructions.


System Requirements :

	Software will run on PC, Linux or Macintosh (tested on Windows7,10, Ubuntu v12 Mac OS X 'Catalina', but anything that will run ImageJ)
	No specific minimum RAM is required but will limit the size of images that can be analysed (Recommend at least 4GB, you will need 1.5GB free
	 to run the demo data)
	A Multi-core processor is also recommended but not essential

Software Requirements :
	You will need a copy of ImageJ of Fiji installed. (tested on Fiji v1.52)
	It should run on any version of ImageJ / Fiji that has java 8.0 or later (java included with ImageJ/Fiji)
	To find the java version in ImageJ/Fiji from the menu select 'Help', 'About ImageJ'
	
Hardware Requirements :
	No non-standard Hardware is required.

***********
Installation
***********
If you have a previous version of the Cox Group Plugin please remove it first.

To install the plugin

Method1:	From the ImageJ/Fiji menu select 'Plugins', 'Install Plugin'. Browse to where you saved this software and select the jar
		file 'CoxGroup_Plugin.jar'

Method2:	just copy the jar file 'CoxGroup_Plugin.jar' to the 'plugins folder' of you ImageJ/Fiji installation.

		On windows10 this should be 'C:\Users\Public\fiji-win64\Fiji.app\plugins'
		On all systems search for 'ImageJ.cfg' should find the ImageJ/Fiji installation folder.

After closing and restarting ImageJ/Fiji the software will appear in the plugins menu.
Installation should only take seconds.

*********
Demo Data
*********

Instructions on use and how to use the example data are included in the html file accessed form the ImageJ/Fiji plugins menu.
	
	for HAWK:	'Plugins', 'Cox Group', 'Hawk', 'Getting Started'
	for HAWKMAN:	'Plugins', 'Cox Group','Hawkman', 'HAWKMAN Help'

Expected output on the example data is included in the help menu

The run time to perform the analysis depends on the number of length scales and the size of the images.
	To analyse the included example data to the default 10 length scales should take ca. 20 seconds on a old/modest PC
	To analyse all 27 length scales up to the PSF will be about 5 minutes.

	Large images at many length scales may take a few tens of minutes and a more powerful PC is advised.
	Progress is displayed in the ImageJ/Fiji log. You can abort at the current length scale by pressing 'Escape'

****************
Trouble Shooting
****************

If ImageJ or FIJI produce an error about insufficient memory to run HAWK or HAWMAN it may be that not enough system RAM has been allocated to ImageJ
As standard this is quite low. It can be increased by selecting 'Exit', 'Options', 'Memory & threads'. Here you can increase the amount of RAM to
be closer to the total SYSTEM ram (eg 90%)
